# Transcriptional characterization of iPSC-derived microglia as a model for therapeutic development in neurodegeneration

**DOI:** 10.1101/2023.03.09.531934

**Authors:** Gokul Ramaswami, Yeliz Yuva-Aydemir, Brynn Akerberg, Bryan Matthews, Jenna Williams, Gabriel Golczer, Jiaqi Huang, Dann Huh, Linda C. Burkly, Sandra J. Engle, Alfica Sehgal, Alla A. Sigova, Robert T. Fremeau, Yuting Liu, David Bumcrot

## Abstract

**Background:** Microglia are the resident immune cells in the brain that play a key role in driving neuroinflammation, a hallmark of neurodegenerative disorders. Inducible microglia-like cells have been developed as an in vitro platform for molecular and therapeutic hypothesis generation and testing. However, there has been no systematic assessment of similarity of these cells to primary human microglia along with their responsiveness to external cues expected of primary cells in the brain.

**Methods:** In this study, we performed transcriptional characterization of commercially available human inducible pluripotent stem cell (iPSC)-derived microglia-like (iMGL) cells by bulk and single cell RNA sequencing to assess their similarity with primary human microglia. To evaluate their stimulation responsiveness, iMGL cells were treated with Liver X Receptor (LXR) pathway agonists and their transcriptional responses characterized by bulk and single cell RNA sequencing.

**Results:** Bulk transcriptome analyses demonstrate that iMGL cells have a similar overall expression profile to freshly isolated human primary microglia and express many key microglial transcription factors and functional and disease-associated genes. Notably, at the single-cell level, iMGL cells exhibit distinct transcriptional subpopulations, representing both homeostatic and activated states present in normal and diseased primary microglia. Treatment of iMGL cells with LXR pathway agonists induces robust transcriptional changes in lipid metabolism and cell cycle at the bulk level. At the single cell level, we observe heterogeneity in responses between cell subpopulations in homeostatic and activated states and deconvolute bulk expression changes into their corresponding single cell states.

**Conclusions:** In summary, our results demonstrate that iMGL cells exhibit a complex transcriptional profile and responsiveness, reminiscent of in vivo microglia, and thus represent a promising model system for therapeutic development in neurodegeneration.

## Background

Microglia are the primary resident immune cells in the brain. Derived from a myeloid lineage, microglia fulfill critical roles in immune surveillance and phagocytosis of cells and debris caused by injury, disease, and aging [1]. Additionally, microglia play important roles in neuronal homeostasis by regulating synaptogenesis and synaptic pruning [2]. Dysregulation of microglial functions are heavily implicated in neurodegenerative disorders including Alzheimer’s disease (AD) [3, 4] and Parkinson’s disease (PD) [5]. Thus, studying the role of microglia in neurodegenerative disorders is essential for developing effective therapies.

Currently, it is untenable to purify sufficient amounts of ex vivo human microglia from brain tissue to perform multifaceted experiments. Additionally, microglia isolated from brain tissue undergo rapid transcriptomic and phenotypic changes when transferred to in vitro conditions [6]. These challenges have facilitated the development of protocols to differentiate iPSCs into iMGL cells [7–12]. These protocols transition iPSCs through hematopoietic precursor cells (HPCs), erythro-myeloid progenitors (EMPs), and finally into iMGL cells in culture [13]. Alternatively, HPCs can be transplanted into early postnatal mouse brain where they develop into microglia [14–16]. These xenotransplant microglia (xMG) more closely resemble the transcriptomic profile of ex vivo microglia than iMGL cells, however they are difficult to produce at large scale.

The goal of this study is to assess commercially available iMGL cells, from Fujifilm Cellular Dynamics, Inc. (Commercial iMGL cells), as a platform for functional studies and target discovery in neurodegeneration. To characterize transcriptional and cellular heterogeneity of Commercial iMGL cells in comparison to primary human microglia, we performed single-cell and bulk RNA sequencing of iMGLs cells and compared results with a number of publicly available transcriptomic data. Then, to assess transcriptional responsiveness of different iMGL cell subpopulations to known neuroprotective agents, we treated Commercial iMGL cells with two LXR pathway agonists [17–19].

## Methods

### Cell Culture

Cell culture was performed as described previously [12] with modifications as described below. iPSC-derived microglia were obtained from FujiFilm Cellular Dynamics, Inc. (clone01279.107, lot 105093). Cells were thawed at 37°C. Each vial was rinsed with 1 mL of culture medium (**Table** 1) and spun at 1,000 x g for 10 minutes. Cells were transferred to 8 mL of culture medium and plated at a density of 5 x 10^4^ cells/cm^2^ in a plate format based on the cell input requirements for a given experiment. Specifically, 6-well format for single-cell and 48-well format for bulk RNA-seq experiments. Cells were grown between 0 and 4 days in culture.

**Table 1.**
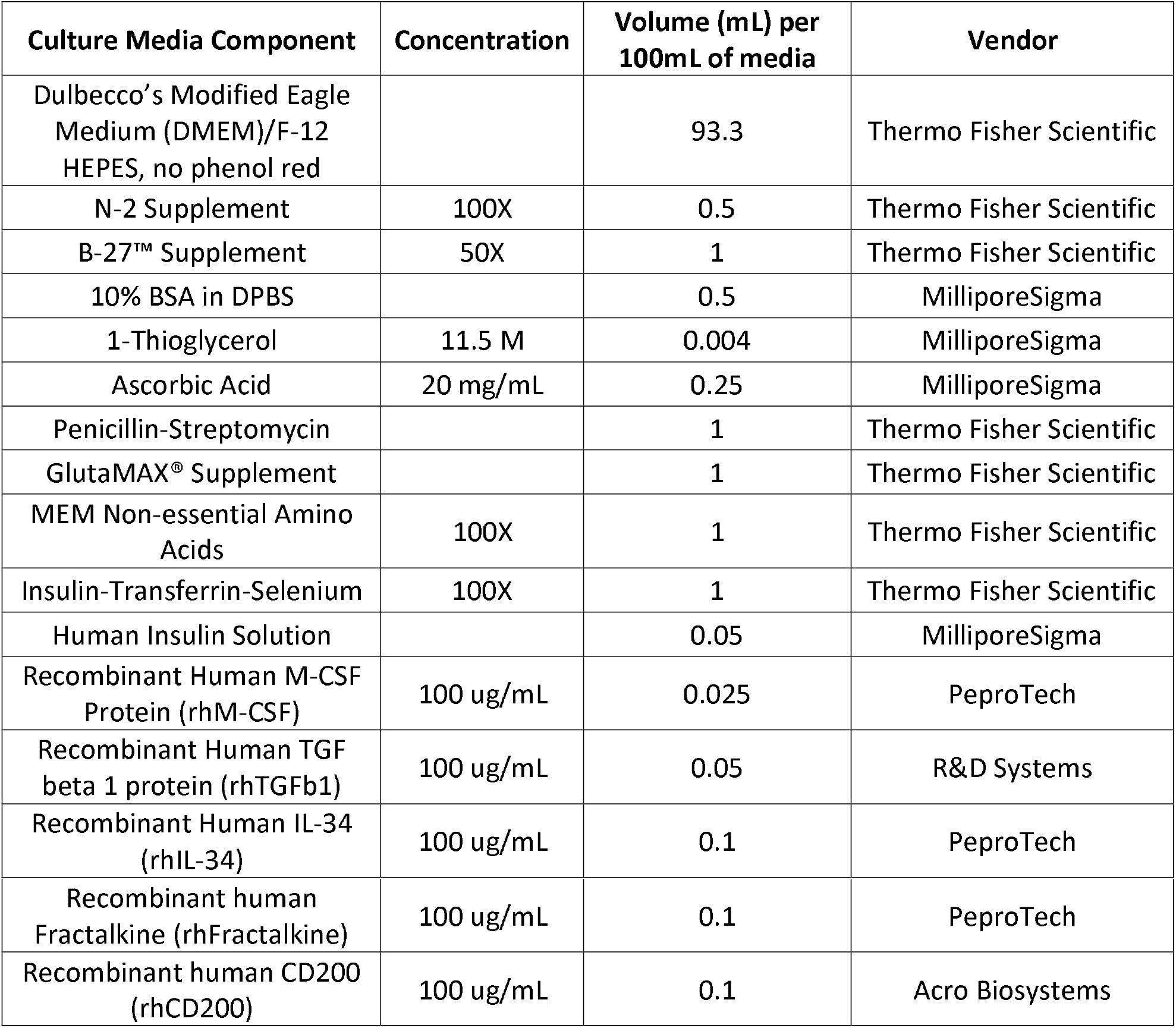
Composition of culture medium

### Immunostaining

Cells were plated on a black glass bottom 96-well plate at a density of ~50,000 cells/cm^2^. On Day 4, cells were washed with 1X phosphate buffered solution (PBS) and fixed with 4% paraformaldehyde in PBS for 15 minutes at room temperature. Permeabilization of the cells was done with 1X PBS containing 0.25% Triton^™^X-100. Following 1 hour blocking in 1X PBS with 5% goat serum and donkey serum at room temperature, goat anti-*IBA1* (Abcam plc, Cat. No. ab5076; 1:100) was added in the blocking solution and incubated at 4°C overnight. The next day, cells were washed 3 times with PBS for 5 min and stained with donkey anti-goat Alexa Fluor^®^ conjugated secondary antibody (Thermo Fisher Scientific Inc., Cat. No. A110-55) at 1:500 for 1 hour at room temperature in the dark. After secondary antibody staining, cells were washed 3 times with PBS and imaged on an Olympus^®^ IX71 microscope (Olympus Corp.).

### RNA Isolation, cDNA synthesis, and bulk RNA sequencing library preparation

For bulk RNA sequencing experiments, medium was removed and cells were lysed directly on plate in RNA lysis buffer and processed using the MagMAX^®^ *mir*Vana Total RNA Isolation Kit (Thermo Fisher Scientific Inc., Cat. No. A27828) according to the manufacturer’s instructions. cDNA was generated from total RNA using Superscript^®^ IV Reverse Transcriptase (Thermo Fisher Scientific Inc., Cat No. 18090050). Bulk RNA-seq library preparation was performed with 500 ng of total RNA using the NEBNext^®^ Poly(A) mRNA Magnetic Isolation Module (New

England Biolabs, Inc., Cat. No. E7490) and NEBNext^®^ Ultra^™^ Directional RNA Library Prep Kit for Illumina^®^ (New England Biolabs, Inc., Cat. No. E7420). All libraries were dual-indexed using 12 cycles of PCR amplification using NEBNext^®^ Multiplex Oligos for Illumina^®^ Dual Index Primer Set (New England Biolabs, Inc., Cat. No. E7600). Library quality control was performed by measuring concentration with Qubit^®^ dsDNA HS Assay Kit (Thermo Fisher Scientific Inc., Cat. No. Q32854) and fragment size distribution with the Agilent^®^ High Sensitivity DNA Kit (Agilent Technologies, Inc., Cat. No. 5067-4626). Libraries were sequenced as paired-end 150×150bp on a NovaSeq^®^ 6000 System (Illumina, Inc.).

### Single Cell RNA-sequencing library preparation

Single cell experiments were performed in parallel with the bulk RNA-seq experiments described above. Cells were first washed with PBS, then trypsinized for 10 minutes at 37°C using TrypLE^®^Express Enzyme (Thermo Fisher Scientific Inc., Cat. No.12604013). Trypsinization was halted by the addition of equal volume of warm medium, cells were spun at 1,000 x g for 10 minutes and resuspended in 0.04% PBS / bovine serum albumin (BSA). Trypan blue was used to assess cell number and viability (>85% in all cases, typically >90%) using a Cellometer^®^ automated cell counter (Nexcelcom Bioscience, LLC). Gel bead-in-emulsion (GEM) encapsulation and single cell indexing reactions were performed using a Chromium^™^ Controller instrument (10x Genomics, Inc., Cat. No. 1000202, version 3.1 chemistry). Single-cell 3’ RNA-seq libraries were prepared using the Chromium^™^ Next GEM Single Cell 3’ GEM, Library and Gel Bead Kit (10X Genomics, Inc., Cat. No. 1000121), according to the manufacturer’s instructions, specifically targeting 2-3,000 cells per replicate and 13 rounds of PCR amplification. Technical replicates were processed independently from trypsinization through sequencing (cells thawed together then plated in independent wells). Libraries were sequenced on a NovaSeq^®^ 6000 System, with paired end 150×150bp sequencing.

### LXR agonist treatment

Cells were plated at 150,000 cells per well on 24-well plates in culture medium. After 24 hours at 37°C, medium was replaced with fresh medium. At day 3, cells were treated with either dimethyl sulfoxide (DMSO) or different doses (30 nM or 100 nM) of T0901317 (MedChemExpress LLC, Cat. No. HY-10626) and GW3569 (30 nM or 300 nM) (MedChem Express LLC, Cat. No. HY-10627A) and incubated for 24 hours. After 24 hours, medium was discarded, and cells were directly lysed on the plate for RNA extraction using the MagMax^®^ *mir*Vana Total RNA Isolation Kit. Bulk and single cell RNA sequencing were performed as described above.

### Time course bulk RNA-seq processing

FastQ files were downloaded from 5 external datasets of microglia related samples [6, 14, 15, 20, 21] (Accession numbers - Gosselin: dbGaP phs001373.v2.p2, Olah: Synapse syn11468526, Galatro: GEO GSE99074, Hasselmann: GEO GSE133432, Svoboda: GEO GSE139194). Samples were removed with less than 10,000,000 reads. FastQ files were processed through a uniform RNA-sequencing pipeline. Briefly, reads were aligned to the hg38 genome with STAR [22] and RSEM was used [23] to quantify the expression of genes in transcripts per million (TPM) using Gencode annotation version 24 [24]. Samples were removed with less than 60% alignment rate after mapping. For external datasets downloaded from literature sources, paired-end samples were removed with greater than 50% duplicate read fraction after mapping. Ribosomal and mitochondrial genes were removed, those that start with “RP” or “MT”. Gene expression values were transformed into log2 (TPM + 0.01) to stabilize the variance. This dataset was used for the assessment of progenitor and monocyte markers. For principal components analyses (PCA) and the assessment of key transcription factors (TFs), marker genes, and disease genes, lowly expressed genes with a median TPM < 1 were removed, leaving us with a dataset of 12,392 genes. ComBat [25] was used to correct for batch effects using the dataset source as the correction factor.

### Principal Components Analysis

The set of all expressed genes was used as features to run PCA. Before running PCA, the expression levels of each gene were scaled across samples by subtracting the mean value and dividing by the standard deviation. PCA was run using the prcomp function in R. For the quantitative comparison in **Figure 1c**, Euclidean distances were calculated between all pairs of samples using their loadings on PC1 and PC2 with the pdist function in R.

**Figure 1.**
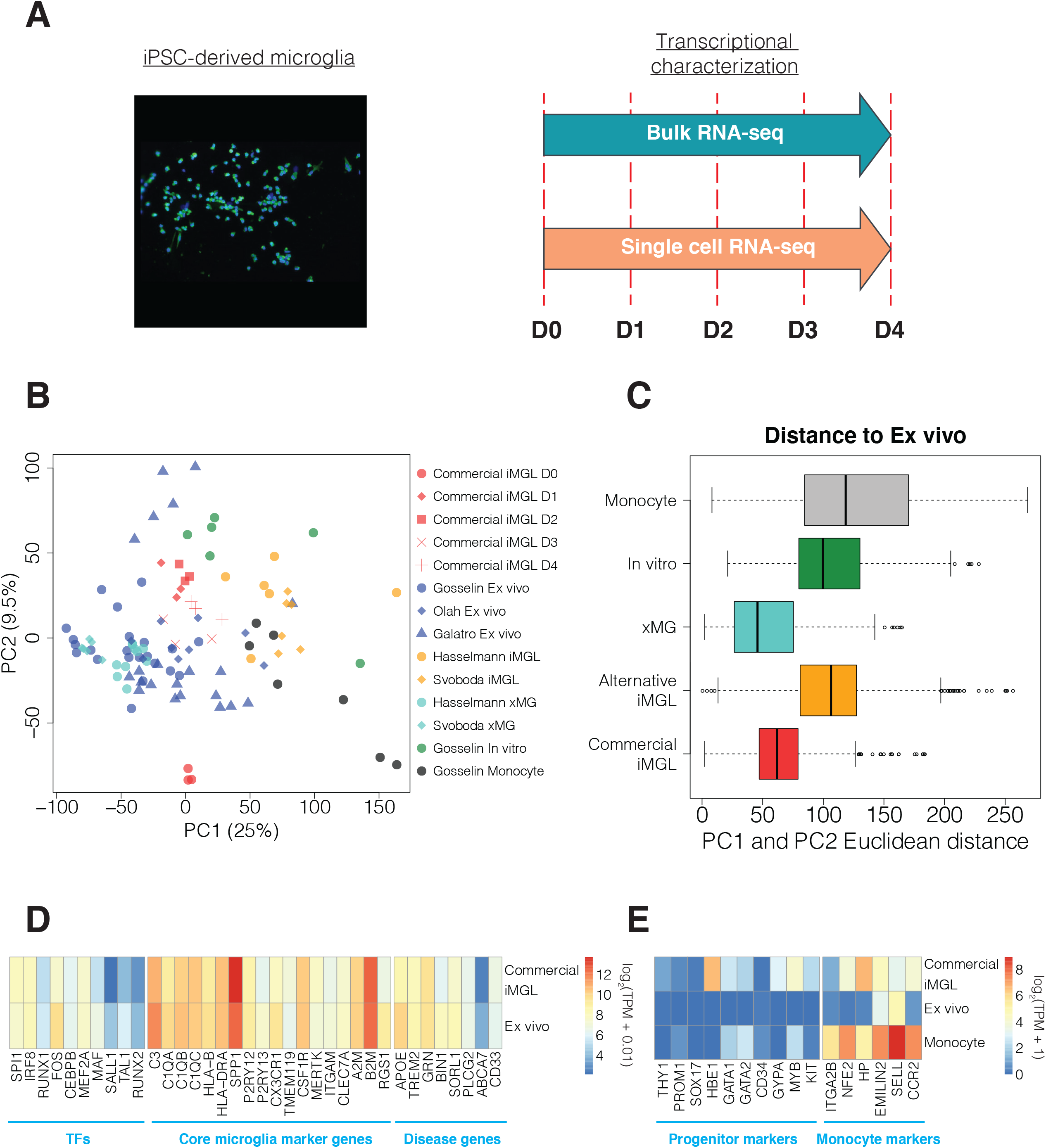
Gene expression comparison between Commercial iMGL cells and microglia-related datasets. **A**) Representative immunofluorescence image of Commercial iMGL cells stained with *IBA1* (green) and DAPI (blue). This study performed bulk and single cell transcriptional profiling of Commercial iMGL cells between 0 and 4 days in culture. **B**) PCA clustering using top 2 PCs for Commercial iMGL cell samples and microglia comparator datasets. PCA was run using all expressed genes. **C**) Euclidean distances for all pairs of samples between ex vivo microglia and each of the other microglia groups. Euclidean distances were calculated using sample loadings on PC1 and PC2. **D**) Comparison of gene expression levels for key microglia TFs, core marker genes, and disease risk genes between Commercial iMGL cells at Days 1-4 and ex vivo microglia. Expression levels displayed are the median values across samples. **E**) Comparison of gene expression levels for HPC and EMP progenitor and monocyte markers between Commercial iMGL cells at Days 1-4, ex vivo microglia, and monocytes. Expression levels displayed are the median values across samples.

### Statistical comparison of cell sources using PCA

For the statistical analysis in **Supplementary Figure 3**, PCA was run separately across all pairs of cell sources. For each independent pair, meaningful principal components (PCs) were identified which explained a greater proportion of variance than random noise. One hundred permutated datasets were created by randomly shuffling sample labels for each gene and the variance explained by the top PC in each permutated dataset was calculated. A threshold for variance explained by meaningful PCs was defined as 2 standard deviations above the mean value for the top component across the 100 permuted datasets. Sample loadings on the meaningful PCs were used to quantitatively access separability for the pair of datasets using 2 methods: SigClust [26] and Silhouette [27]. The SigClust score was calculated using the sigclust package in R. The Silhouette score was calculated using the cluster package in R. One hundred permuted datasets were generated by shuffling the sample labels for the meaningful PCs and re-running the 2 methods. For each method, an empirical p-value was calculated to assess the significance of dataset separability by identifying the proportion of permutated datasets with a larger separability score then the actual score. In the case of Sigclust, this was the proportion of permutated datasets with a lower score, because lower SigClust scores correspond to greater separability. For Silhouette, this was the proportion of permutated datasets with a higher score, because higher Silhouette scores correspond to greater separability.

### Time course single cell RNA-seq processing

FastQ files for each single cell RNA-seq sample were processed using Cell Ranger (10x Genomics, Inc.) to generate count matrices of genes per cell. Count matrices were processed for downstream analyses using Seurat version 3.2 [28]. Low quality cells were removed, potentially empty droplets or doublets by imposing the following filtering criteria for each cell: number of genes detected > 1500, number of reads per cell < 100,000, and mitochondrial RNA (mtRNA) fraction < 20%. We also filtered any genes that were detected in less than 3 individual cells. This left us with a dataset of 18,426 genes across 31,984 cells collectively. Gene expression values were log-transformed using the NormalizeData function and the top 2,000 variable genes were identified using the FindVariableFeatures function. Expression levels for each gene were scaled and centered using the ScaleData function. Louvain clustering was performed with the top 20 principal components using the FindNeighbors and FindClusters functions with a resolution parameter of 0.4. For visualization, the variance across the top 20 principal components was reduced into 2 Uniform Manifold Approximation Projection (UMAP) dimensions using the RunUMAP function. For cells in each cluster, preferentially expressed marker genes were identified using the FindAllMarkers function with the following criteria: adjusted Wilcox p-value < 0.01, detected percentage > 40%, and Log2(fold-change) > 0.6. GO enrichment analyses was performed for the marker gene sets using the clusterProfiler package in R [29].

### LXR agonist treatment bulk RNA-seq processing

FastQ files were processed as described above. Differential gene expression analysis was run using the DESeq2 package in R [30]. The DESeq function was used to perform median of ratios normalization of the count data. Four contrasts were run: GW3569 30nM vs DMSO, GW 300 nM vs DMSO, T0901317 30 nM vs DMSO, and T0901317 100 nM vs DMSO. To identify DEGs, the following criteria were used: absolute value of log_2_ fold change > 0.6, adjusted p-value < 0.05, and the maximum TPM value across samples > 1. The enrichR package in R [31] was used to run pathway enrichment analysis. For each contrast, both up- and downregulated DEGs were mixed and run against public databases including KEGG_2019_Human, GO_Biological_Process_2018 and Reactome_2016. The significance of the enrichment and overlapping genes were generated by the enrichr function for each pathway. Significantly enriched pathways were identified with an adjusted p-value < 0.05.

### LXR agonist treatment single cell RNA-seq processing

FastQ files were processed as described above. After filtering of low quality cells, we started with a dataset of 19,517 genes in 43,312 cells, collectively. Data normalization and clustering were performed as described above, expect we used a resolution parameter of 0.2 for Louvain clustering. For each of the 4 contrasts, DEGs were identified in each cluster using the FindAllMarkers function with the following criteria: absolute value of log_2_ fold change > 0.6, adjusted Wilcox p-value < 0.05, and percentage of detected cells > 10%.

## Results

### Comparison of Commercial iMGL cell bulk transcriptome data with publicly available primary microglia-related datasets

*We* verified expression of microglia marker *IBA1 on* Commercial iMGL cells by immunostaining (**Figure 1a**). Bulk RNA sequencing was performed on cells grown between 0 and 4 days in culture (**Figure 1a and Methods**). As comparator datasets, we collected published bulk RNA sequencing data from five studies [6, 14, 15, 20, 21], encompassing three human primary microglia datasets (ex *vivo*), two alternative iMGL cell datasets, two xMG datasets, one *in-vitro* cultured human primary microglia dataset, and one human ex vivo monocyte dataset (**Supplementary Table** 1). For all RNA sequencing samples, we started with FastQ files, processed them using a uniform pipeline, and performed batch correction to minimize sources of technical variation between datasets (**Methods and Supplementary Figure** 1).

To assess genome-wide patterns in gene expression, we conducted principal components analysis (PCA) with all expressed genes. We found that xMG and Commercial iMGL cell samples are more closely related to ex vivo microglia than alternative iMGL cells, in vitro cultured microglia, and monocytes by visual (**Figure 1b**) and quantitative (**Figure 1c**) comparisons using the top two principal components (PCs). We also examined two different similarity metrics, SigClust and Silhouette index, with all PCs above noise, which has been determined by permutation tests (**Methods**). The overall similarity was observed in both metrics used (**Supplementary Figure 2**). These results demonstrate that the overall expression profile is similar between ex vivo microglia and Commercial iMGL cells.

Next, we compared the expression of key microglia TFs [32], core marker genes associated with microglia functions [33], and AD risk genes [34] between the datasets. We found the expression of most TFs, marker genes, and disease genes to be similar between ex vivo microglia and Commercial iMGL cells (**Figure 1d and Supplementary Figure 3**). However, there are differences including lower expression of *SALL1* and higher expression of *SPP1* in Commercial iMGL cells, two genes that exhibit expression changes in models of neurodegenerative disease [35, 36]. We also assessed the expression of iPSC, HPC, and EMP marker genes as well as monocyte enriched marker genes [37] (**Figure 1e and Supplementary Figure 3**). We found moderate expression of monocyte markers and multiple progenitor markers including *HBE1, GATA1, GATA2, GYPA, MYB*, and *KIT*, suggesting the presence of a progenitor subpopulation within the Commercial iMGL cell culture. Taken together, these results demonstrate that Commercial iMGL cells largely recapitulate the basal transcriptional profile of ex vivo microglia.

### Single cell transcriptomics identifies multiple subpopulations of cells within Commercial iMGL cells

Microglia exhibit a variety of transcriptional states in response to the local environment [32, 38]. These states are indicative of individual microglia performing different roles including surveillance and neuronal homeostasis (homeostatic microglia) as well as inflammatory response and phagocytosis (activated microglia). To characterize the heterogeneity of Commercial iMGL cells in culture, we performed single-cell RNA sequencing on cells grown between 0 and 4 days in culture (**Figure 1a and Methods**). After removing low quality cells (**Supplementary Figure 4**), we generated a dataset of 31,984 cells. Using de novo clustering, we identified 11 clusters of cells within Commercial iMGL cell cultures (**Figure 2a**). To assign functional states for each cluster, we assessed the expression patterns of genes, from a single cell study of human primary microglia, that mark important aspects of microglial biology including antigen presentation, complement pathway, immune activation, and cell cycle [38] (**Figure 2c and Supplementary Figure** 5). Additionally, we generated gene ontology enrichments of differentially expressed cluster marker genes (**Methods, Supplementary Table 2, and Supplementary Figure 6**). We identified 4 clusters, C1, C3, C7, and C10, indicative of homeostatic microglia with high expression of genes involved in antigen presentation and complement pathways. We identified 5 clusters, C2, C4, C5, C8, and C9, indicative of activated microglia with high expression of immune related genes. Among activated microglia, 2 clusters, C2 and C5, have high expression of immediate-early genes which are rapidly induced upon stimulation [39]. Among homeostatic and activated microglia, we identified 3 clusters of proliferating cells, C3, C7, and C9 with high expression of cell cycle genes. In concordance with bulk expression patterns (**Figure 1e**), we identified 1 cluster, C11, corresponding to myeloid progenitors (**Supplementary Figure 7**). Reassuringly, the proportion of total cells within C11 is less than one percent (**Figure 2b**), demonstrating the low prevalence of incompletely differentiated cells. Of note, AD risk genes were differentially expressed across clusters (**Supplementary Figure 8**), supporting the roles of multiple microglia functions and pathways in the etiology of AD [40].

**Figure 2.**
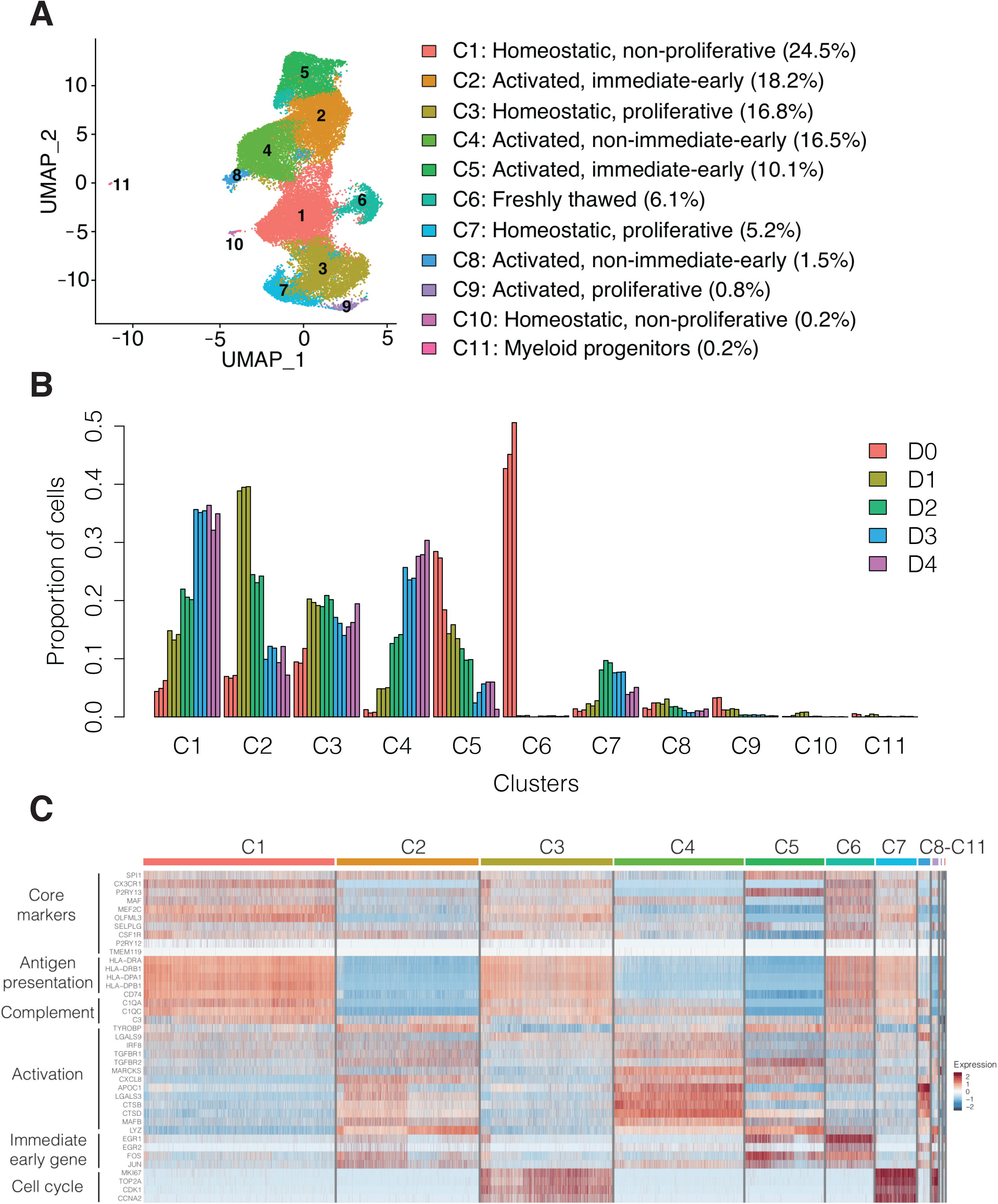
Single cell RNA-seq profiling of Commercial iMGL cell cultures. **A**) Identification of 11 cellular clusters in Commercial iMGL cell cultures. Functional annotations were assigned based on the expression of microglia functional marker genes as described in the **Methods** and shown in **Figure 2c**. **B**) Proportion of cells in each cluster in the time course single cell RNA-seq dataset. **C**) Gene expression heatmap of selected microglia functional marker genes in the time course single cell RNA-seq dataset. Heatmap with a more comprehensive list of marker genes is shown in **Supplementary Figure 5**.

The proportion of cells within each cluster was dynamic over the 4-day time course (**Figure 2b**). One cluster, C6, with high expression of homeostatic and activated marker genes (**Figure 2c**), was only present at D0 and indicative of freshly thawed cells. The proportion of homeostatic microglia in cluster 1, those not expressing proliferating markers, increased over time in culture from 15% of cells at day 1 to 35% of cells at day 4. In contrast, the proportion of proliferative homeostatic microglia in cluster C3 was relatively stable between 1 and 4 days in culture. The proportion of activated microglia in cluster C4, those not expressing immediate-early genes, increased over time in culture from 5% at day 1 to 30% at day 4, while the proportion of activated microglia expressing immediate-early genes in cluster C2 decreased over time in culture from 40% at day 1 to 10% at day 4. Overall, the proportion of microglia subpopulations changes substantially over time in culture, approaching steady state between days 3 and 4.

### Robust bulk transcriptional response of Commercial iMGL cells to LXR pathway agonists

To assess the transcriptional responsiveness of Commercial iMGL cells, we selected two structurally distinct LXR pathway agonists: T0901317 and GW3569 (**Figure 3a**) [41, 42]. LXRs are lipid responsive TFs that form heterodimers with retinoid X receptors (RXRs) to regulate expression of key cholesterol homeostatic genes involved in AD [43]. LXR agonists have previously been shown to reduce amyloid plaque burden, ameliorate neuroinflammation, and improve memory in preclinical AD models [17–19].

**Figure 3.**
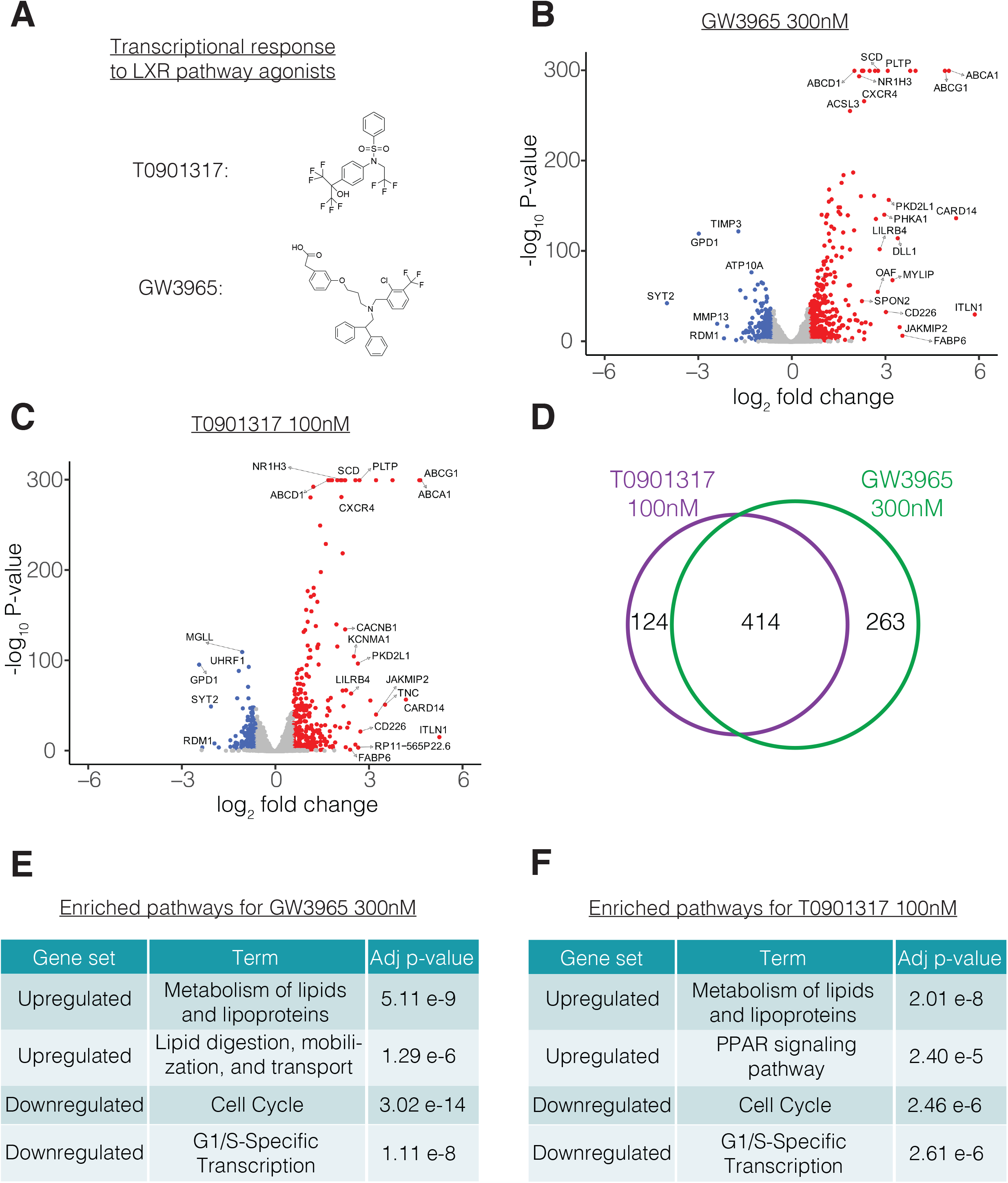
Bulk transcriptional responses of Commercial iMGL cells to LXR agonist treatment. **A**) Chemical structures of LXR pathway agonists GW3965 and T0901317. **B**) Volcano plot showing relationship between expression fold change and significance for differential gene expression between GW3965 300nM and DMSO treated cells. Genes with significant upregulation and downregulation in GW3965 300nM are colored red and blue, respectively. **C**) Volcano plot showing relationship between expression fold change and significance for differential gene expression between T0901317 100nM and DMSO treated cells. Genes with significant upregulation and downregulation in T0901317 100nM are colored red and blue, respectively. **D**) Overlap in DEGs between GW3965 300nM and T0901317 100nM treatments. **E**) Selected pathway enrichments for DEGs with GW3965 300nM treatment. **F**) Selected pathway enrichments for DEGs with T0901317 100nM treatment.

We treated Commercial iMGL cells with two doses of each LXR agonist starting on day 3 of in vitro culture and performed bulk RNA sequencing to assess their transcriptional responses 24 hours later (**Methods and Supplementary Table 3**). We found a marked increase in expression of the established LXR pathway target genes [44] *ABCA1, ABCG1*, and *APOE* (**Supplementary Figure 9**). We systematically identified differentially expressed genes (DEGs) for each treatment (**Methods**). We identified 296, 677, 227, and 538 DEGs for GW3965 30nM, GW3965 300nM, T0901317 30nM, and T0901317 100nM, respectively (**Supplementary Table 4**). In all four stimulus conditions, *ABCA1* and *ABCG1* were among the most highly upregulated genes, as expected with stimulation of the LXR pathway (**Figure 3b-c and Supplementary Figure 10**). Additionally, for both compounds, the expression level of DEGs was dose-dependent, and almost all the DEGs identified at the lower dose were also identified at the higher dose (**Supplementary Figure 11**). Comparing between the two compounds, we found highly overlapping lists of DEGs (**Figure 3d**), suggesting that both compounds are modulating the same cellular pathways. We ran gene ontology and pathway enrichments for the DEGs identified in each of the 4 stimulus conditions (**Supplementary Tables 5-8**). We found similar pathway enrichments in both compounds with upregulated genes enriched in lipid metabolism processes and downregulated genes enriched in cell cycle, extracellular matrix, and inflammatory processes (**Figure 3e-f**). Overall, Commercial iMGL cells exhibited robust bulk transcriptional responses to LXR agonists.

### Differential transcriptional responses to LXR pathway agonists across Commercial iMGL cell subpopulations

To assess cell-to-cell heterogeneity in transcriptional responses of Commercial iMGL cells to LXR pathway agonists, we performed single cell RNA sequencing (**Methods**). After removing low quality cells (**Supplementary Figure 12**), we generated a dataset of 43,312 cells. Using Louvain clustering, we identified 6 clusters of cells (**Figure 4a**) and matched their identities to the previously identified Commercial iMGL cell subpopulations (**Figure 2a**) using a set of microglia functional marker genes [38] (**Supplementary Figure 13**). In general, the proportion of cells within each cluster is consistent between untreated and treated samples (**Figure 4b**). There is a slight reduction in the proportion of proliferative homeostatic cells in cluster C3 which is consistent with the finding that downregulated genes in bulk RNA-seq are enriched within the cell cycle pathway (**Figure 3e-f**).

**Figure 4.**
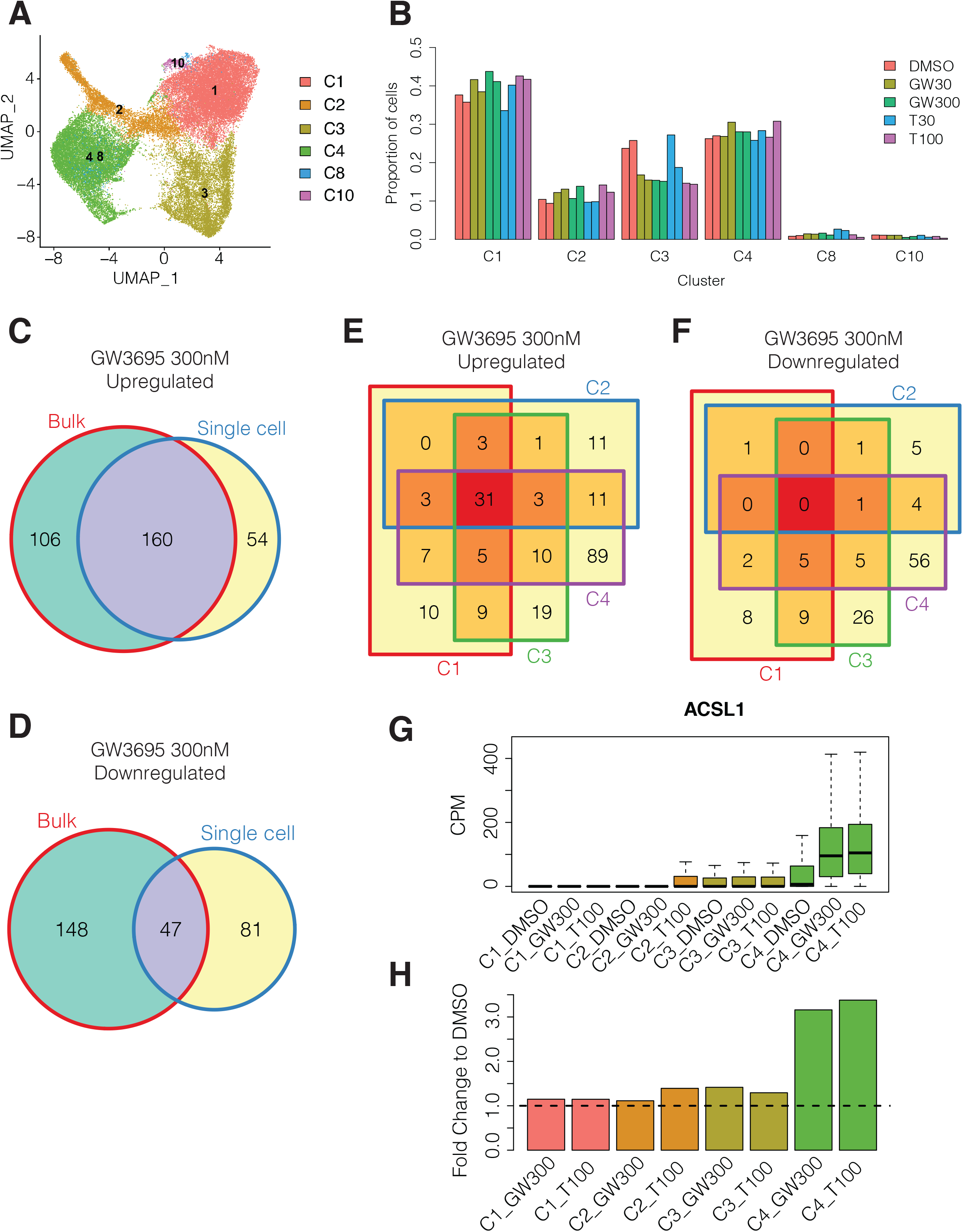
Single cell transcriptional responses of Commercial iMGL cells to LXR agonist treatment. **A**) Identification of 6 single cell clusters in LXR agonist treated Commercial iMGL cells. Cluster names were matched to the time course single cell RNA-seq dataset (**Figure 2a**) using microglia functional marker genes (**Supplementary Figure 13**) as described in the Methods. **B**) Proportion of cells belonging to each single cell cluster in the LXR agonist treatment single cell RNA-seq dataset. **C**) Overlap in upregulated DEGs between bulk RNA-seq and the union of single cell RNA-seq DEGs in clusters C1, C2, C3, C4, C8, and C10 with GW3965 300nM treatment. Bulk DEGs were filtered to genes tested in at least one single cell contrast. **D**) Overlap in downregulated DEGs between bulk RNA-seq and the union of single cell RNA-seq DEGs in clusters C1, C2, C3, C4, C8, and C10 with GW3965 300nM treatment. Bulk DEGs were filtered to genes tested in at least one single cell contrast. **E**) Overlap in upregulated DEGs for single cell clusters C1, C2, C3, and C4 with GW3965 300nM treatment. **F**) Overlap in downregulated DEGs for single cell clusters C1, C2, C3, and C4 with GW3965 300nM treatment. **G**) Expression of *ACSL1* gene in single cells assigned to clusters C1, C2, C3, and C4 from DMSO, GW3965 300nM (GW300), and T0901317 100nM (T100) treatments. **H**) Fold change of *ACSL1* gene compared to DMSO in single cells assigned to clusters C1, C2, C3, and C4 from GW3965 300nM (GW300) and T0901317 100nM (T100) treatments.

We identified LXR agonist-induced DEGs for each cluster (**Methods, Supplementary Figure 14, and Supplementary Table 9**). First, we compared the DEG responses between high and low doses for each treatment within the same cluster. In general, fold change differences correlate with dose for each treatment (**Supplementary Figure 15**). We noticed that clusters C1, C3, and C4 show higher correlations between doses than cluster C2, C8, and C10, for both treatments, suggesting that Commercial iMGL cell subpopulations have differing sensitivities to LXR agonists. Comparing between the two compounds, we found highly overlapping lists of DEGs within each cluster (**Supplementary Figure 16**), and similar gene ontology and pathway enrichments (**Supplementary Table 10**).

Next, we compared the set of DEGs between bulk and single cell RNA-seq. In general, the gene expression changes detected by bulk and single cell analysis are quite similar (cor = 0.7 - 0.81). However, there are many genes that exhibit differences in fold-changes between the two technologies (**Supplementary Figure 17**). The number of DEGs is greater in bulk as compared to single cell, however there is substantial overlap between the two methods (**Figure 4c-d and Supplementary Figure 18**). We found that DEGs detected by bulk only exhibit lower expression levels in the single cell RNA-seq dataset when compared to DEGs detected by both methods (**Supplementary Figure 19**). This finding is consistent with previous reports showing that identification of lowly expressed DEGs by single cell analysis is challenging [45].

Finally, we compared the DEG responses for the high dose of each compound between the four major clusters C1, C2, C3, and C4. Between clusters, the gene expression responses were positively correlated, however the fold change extents were different (**Supplementary Figure 20**). We found that the transcriptional response in cluster 1 was more comparable to cluster 3 (cor = 0.74-0.77) than cluster 2 and cluster 4 (cor = 0.52-0.64), which support our previous characterization of C1/C3 and C2/C4 as representing homeostatic and activated microglia, respectively. We compared the set of DEGs across different clusters in each treatment and found substantial differences in DEGs across clusters, with a particular emphasis on a large number that are specific to cluster C4 after GW3965 300 **nM (Figure 4e-f**) and T0901317 100 nM treatments (**Supplementary Figure 21**). One example of a gene with differential transcriptional responses across clusters is *ACSL1*. The expression of *ACSL1* is upregulated in cluster C4 (FC = 3.16 – 3.38), in response to both compounds, whereas its expression is unchanged in clusters C1, C2 and C3 (FC = 1.11 – 1.42) (**Figure 4g-h**). *ACSL1* is significantly upregulated in bulk RNA-seq (FC = 2.6 – 2.75) (**Supplementary Table 4**), and the single cell dataset demonstrates that this bulk signal is predominantly driven by activated microglia cells in cluster C4. Similarly, we sorted the full list of bulk DEGs into their corresponding single cell clusters (**Supplementary Table 11**). These findings demonstrate that different subpopulations of iMGL cells exhibit heterogeneous transcriptional responses to LXR agonists.

## Discussion

Microglia play important roles in the pathogenesis of neurodegenerative disorders. A substantial increase in neuroinflammation mediated by microglial activation and proliferation is a hallmark of late stage neurodegenerative disorders including AD, PD, and frontotemporal dementia [46]. Two of the most prominent genetic risk factors for late onset AD, *APOE* and *TREM2*, are highly expressed in microglia and are important regulators of cholesterol metabolism and transport in the brain [47]. Therefore, in vitro models, such as iMGL cells, are essential tools for preclinical development. Multiple studies have shown that iMGL cells have transcriptional profiles that are highly similar to ex vivo microglia [12, 15, 48–52]. In this study, we find that iMGL cells, from Fujifilm Cellular Dynamics, Inc., are an attractive commercial option.

Currently, multiple therapeutics targeting APOE and TREM2 in microglia are in development for AD [53–55], including LXR pathway agonists [56]. This is the first study to perform a comprehensive bulk and single cell transcriptional characterization of LXR agonists on iMGL cells. At the bulk level, we find an upregulation of lipid metabolism pathways, consistent with previous findings that positive effects of LXR agonists on AD pathology in preclinical models are largely due to the induction of *ABCA1* expression, which promotes cholesterol clearance and APOE lipidation [17, 47]. Interestingly, at the single cell level, we identified heterogeneous responses across subpopulations of iMGL cells including a prominent response in activated microglia. We found a downregulation of inflammatory response pathways, consistent with a previous report that treatment of mouse microglia-derived cell line BV2 with GW3569 causes an attenuation of neuroinflammation [57].

Although Commercial iMGL cells show promising potential, they do not completely recapitulate all aspects of microglial biology. The transcriptional factor *SALL1*, an important determinant of microglia identity [58], is not expressed in Commercial iMGL cells (**Figure 1d and Supplementary Figure 3**). However, xenotransplant microglia exhibit expression of *SALL1*, indicating that iMGL cells have the potential to turn on the *SALL1*-mediated transcriptional program (**Supplementary Figure 3**). Moving forward, optimization of differentiation protocols and culture media components as well as further methods development, including direct cell conversion [48], will continue to make iMGL cells an attractive model system for therapeutic development in neurodegeneration.

## Conclusions

In this study, we performed comprehensive bulk and single cell transcriptional characterization of Commercial iMGL cells. We find that Commercial iMGL cells closely resemble ex vivo microglia at the overall transcriptome level and express most, but not all, key microglia marker genes. We identified 11 subpopulations of cells representing homeostatic, activated, and proliferating states, replicating the heterogeneity of ex vivo microglia [38]. Commercial iMGL cells stabilize after 3 days in culture and we treated them with two distinct LXR pathway agonists to assess their transcriptional responsiveness. At the bulk level, Commercial iMGL cells respond by upregulation of lipid metabolism pathways and downregulation of cell cycle pathways. At the single cell level, the transcriptional responses differ between homeostatic and activated microglia. Overall, our results demonstrate that Commercial iMGL cells exhibit a basal transcriptional profile, cellular heterogeneity, and transcriptional plasticity that is comparable to in vivo microglia.

## List of Abbreviations

iPSC: induced pluripotent stem cells
iMGL cells: iPSC-derived microglia-like cells
LXR: liver X receptor
xMG: xenotransplant microglia
PCA: principal components analysis
DEG: differentially expressed gene
DMSO: dimethyl sulfoxide
HPC: hematopoietic precursor cells
EMP: erythro-myeloid progenitors
AD: Alzheimer’s disease
PD: Parkinson’s disease
TF: transcription factor

## Declarations

### Ethics approval and consent to participate

Not applicable

## Consent for publication

Not applicable

## Availability of data and materials

Raw and processed data from bulk and single cell RNA sequencing experiments have been deposited in the NCBI Gene Expression Omnibus (GEO) under accession number GSE226081.

## Authors’ contributions

G.R. performed computational analyses and wrote the manuscript draft. Y.Y., B.A., B.M., and J.W. performed experiments. G.G., J.H., and Y.L. performed computational analyses. D.H., L.C.B., S.J.E., A.S., A.A.S., R.T.F., Y.L., and D.B. supervised the experiments and computational analyses. All authors aided in the interpretation of results; read, reviewed, and approved the manuscript for publication.

## Acknowledgements

The results published here are in part based on data obtained from the AD Knowledge Portal (https://adknowledgeportal.org). The results published here are in part based on data obtained from dbGaP (PI: Glass, Institute: NINDS).

## Additional files

Additional file 1: Supplementary_Figures_v5.pdf

Additional file 2: STable1.xlsx

Additional file 3: STable2.xlsx

Additional file 4: STable3-8.xlsx

Additional file 5: STable9.xlsx

Additional file 6: STable10.xlsx

Additional file 7: STable11.xlsx

