## Supplementary Figures for "Transcriptional characterization of iPSC-derived microglia as a model for therapeutic development in neurodegeneration"

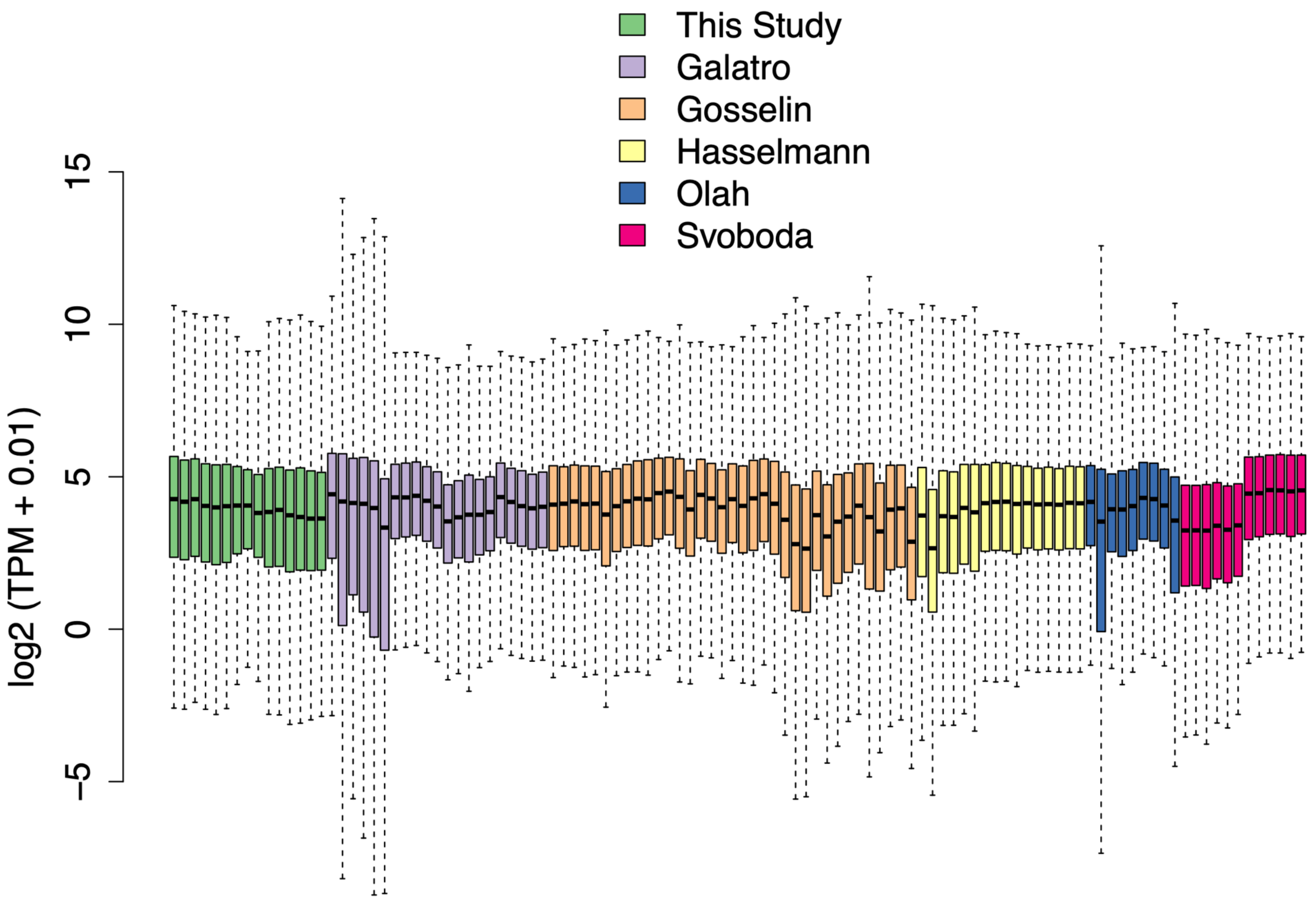
This Study
Galatro
Gosselin
Hasselmann
Olah
Svoboda
log2 (TPM + 0.01)
15
10
5
0
−5

**Supplementary Figure 1.** Distribution of gene expression values for 12,392 expressed genes across the Commercial iMGL time course and external bulk RNA-seq datasets.

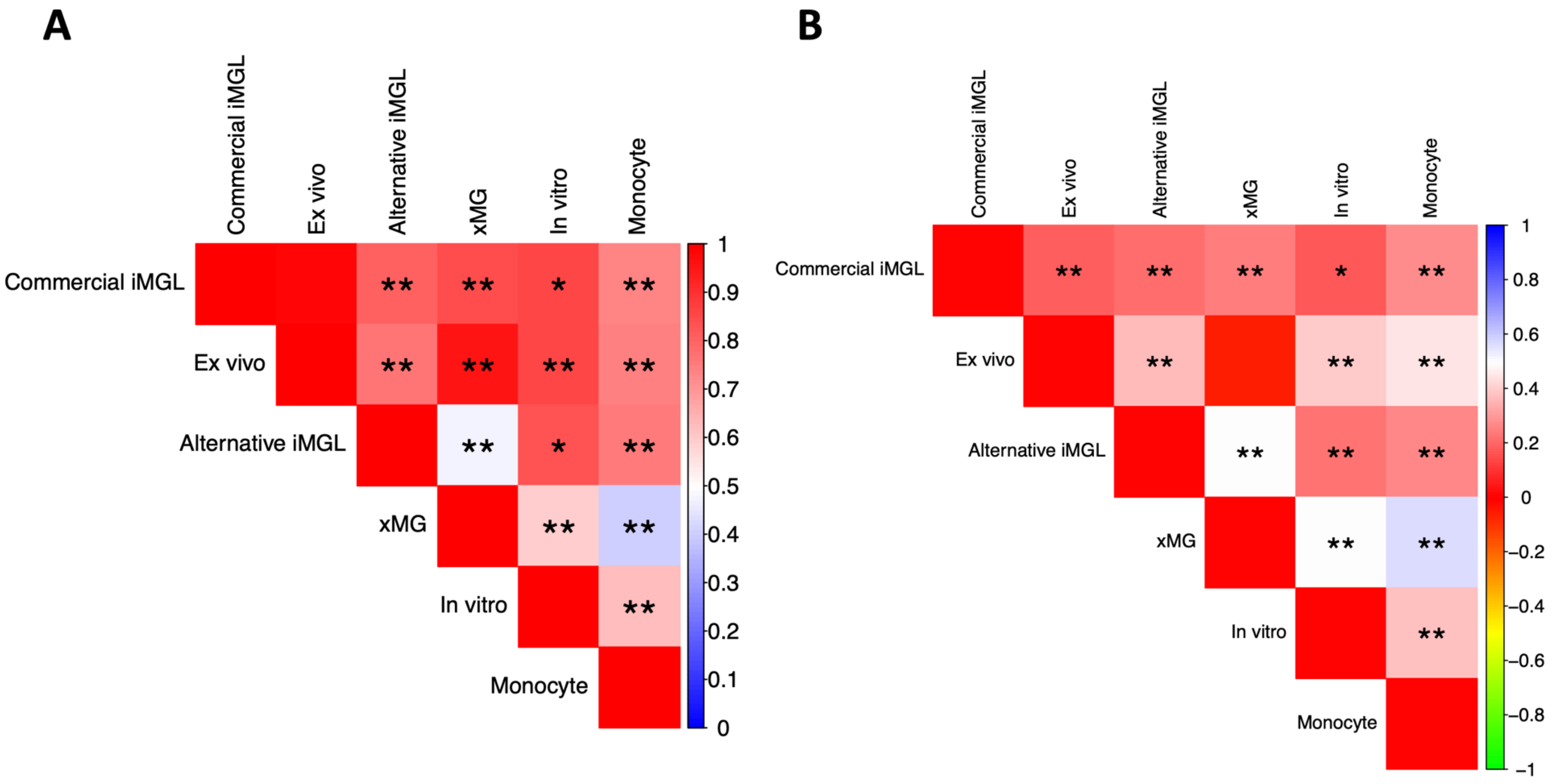

A
Commercial iMGL
Ex vivo
Alternative iMGL
xMG
In vitro
Monocyte
** ** * **
** ** ** **
** * **
** **
**
1
0.9
0.8
0.7
0.6
0.5
0.4
0.3
0.2
0.1
0
B
Commercial iMGL
Ex vivo
Alternative iMGL
xMG
In vitro
Monocyte
** ** ** * **
** ** **
** ** **
** **
**
1
0.8
0.6
0.4
0.2
0
−0.2
−0.4
−0.6
−0.8
−1

**Supplementary Figure 2.** Quantitative PCA comparisons

**A**) Pairwise Sigclust scores comparing Commercial iMGL samples and microglia comparator datasets using principal component analysis on all expressed genes. Each cell represents a comparison between two cell systems and is colored by the Sigclust score (0 to 1). * empirical p-value < 0.05, ** empirical p-value < 0.01

**B**) Pairwise average Silhouette scores comparing Commercial iMGL samples and microglia comparator datasets using principal component analysis on all expressed genes. Each cell represents a comparison between two cell systems and is colored by the average Silhouette score (-1 to 1). * empirical p-value < 0.05, ** empirical p-value < 0.01

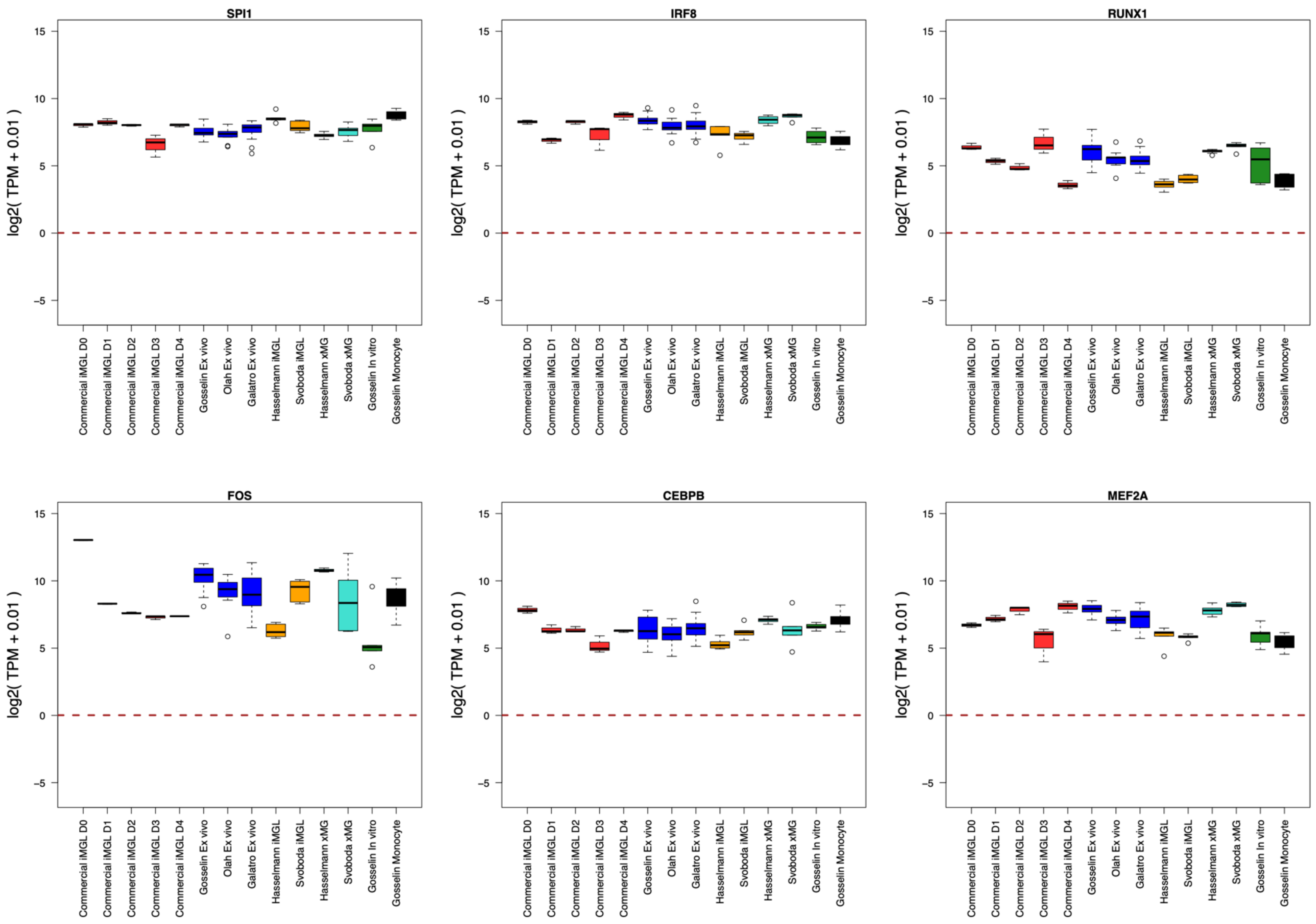

SPI1
IRF8
RUNX1
FOS
CEBPB
MEF2A
log2( TPM + 0.01 )
15
10
5
0
−5
Commercial iMGL D0
Commercial iMGL D1
Commercial iMGL D2
Commercial iMGL D3
Commercial iMGL D4
Gosselin Ex vivo
Olah Ex vivo
Galatro Ex vivo
Hasselmann iMGL
Svoboda iMGL
Hasselmann xMG
Svoboda xMG
Gosselin In vitro
Gosselin Monocyte

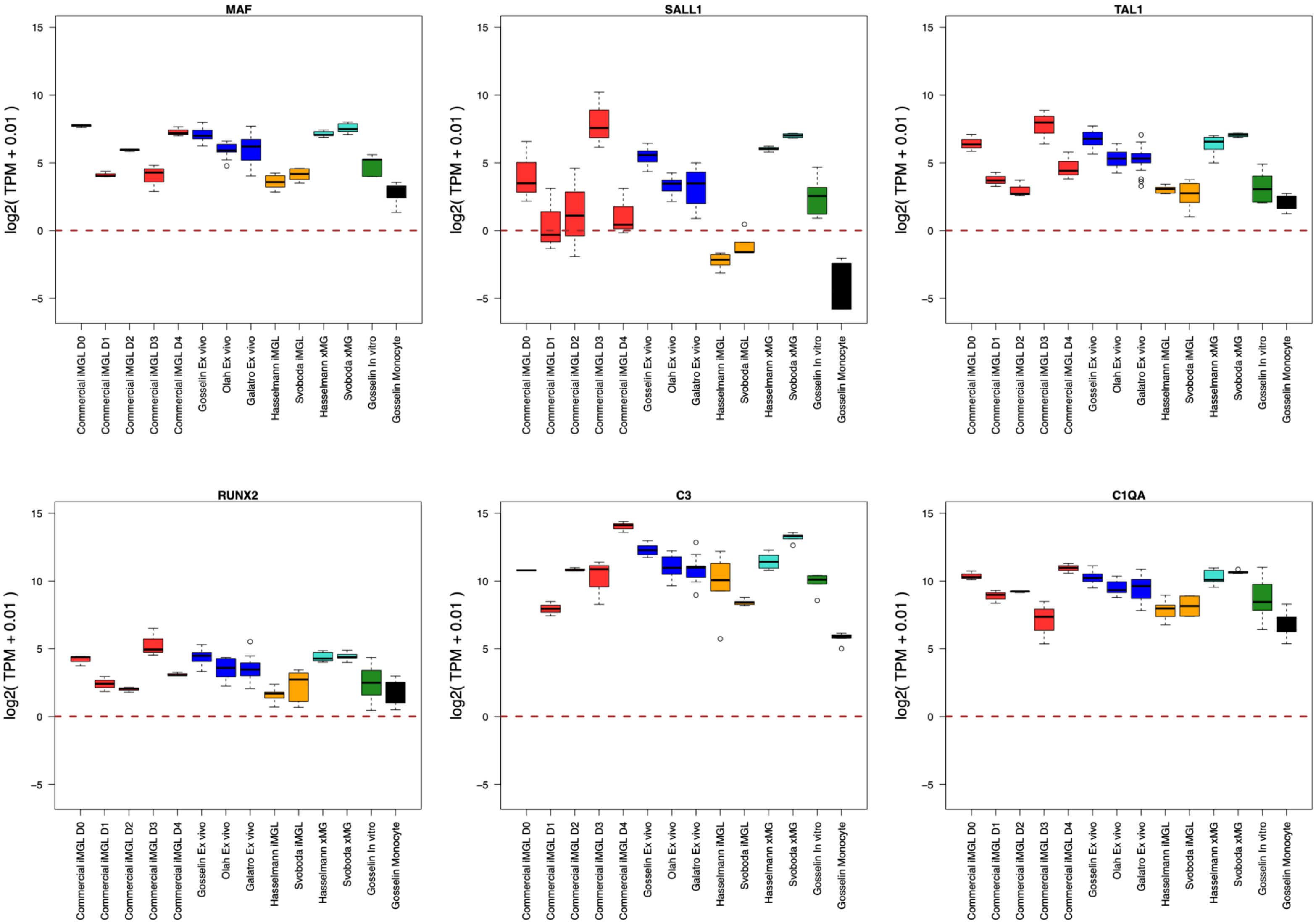
MAF
SALL1
TAL1
RUNX2
C3
C1QA
log2( TPM + 0.01 )
15
10
5
0
−5
Commercial iMGL D0
Commercial iMGL D1
Commercial iMGL D2
Commercial iMGL D3
Commercial iMGL D4
Gosselin Ex vivo
Olah Ex vivo
Galatro Ex vivo
Hasselmann iMGL
Svoboda iMGL
Hasselmann xMG
Svoboda xMG
Gosselin In vitro
Gosselin Monocyte

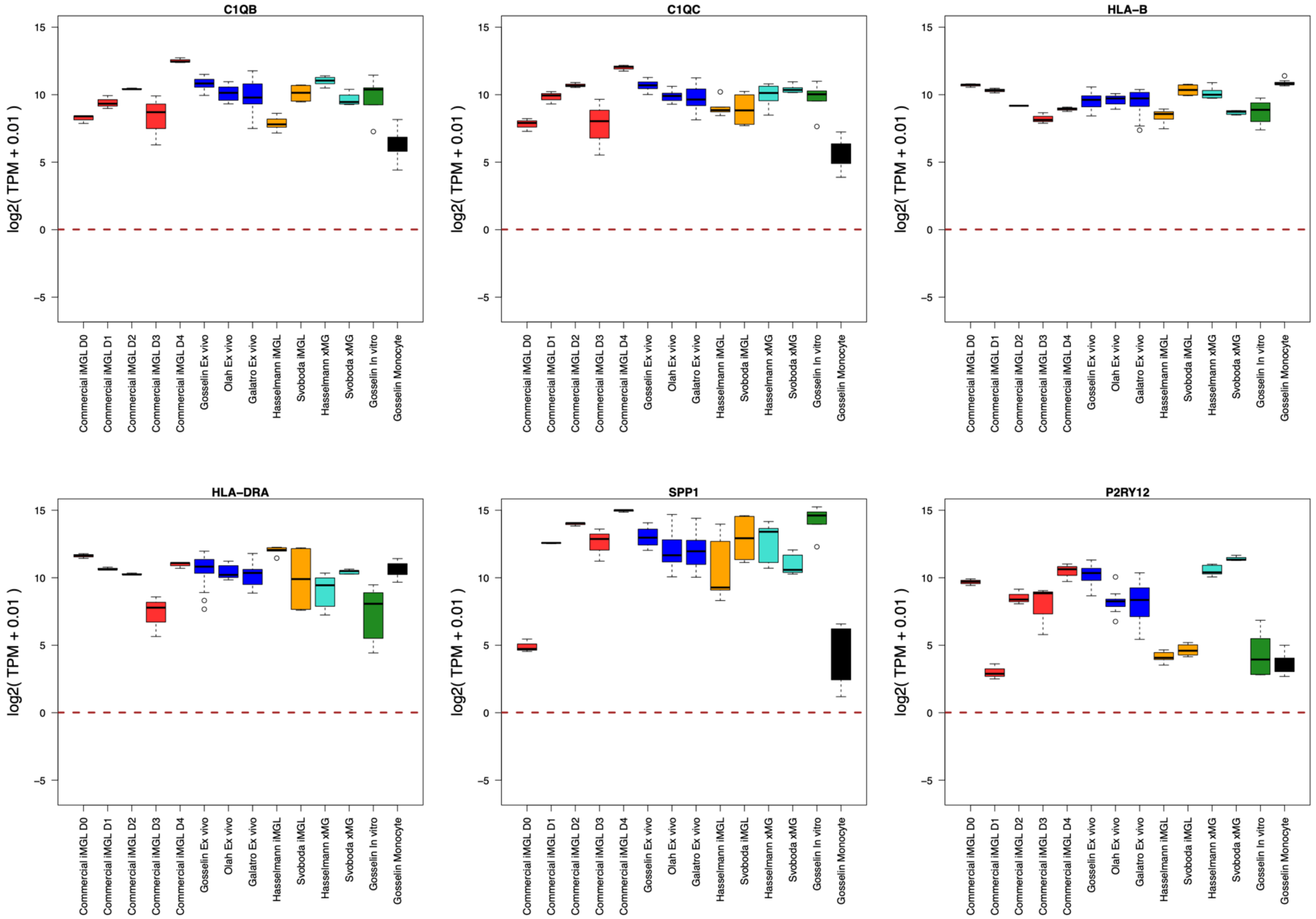
C1QB
C1QC
HLA−B
HLA−DRA
SPP1
P2RY12
log2( TPM + 0.01 )
15
10
5
0
−5
Commercial iMGL D0
Commercial iMGL D1
Commercial iMGL D2
Commercial iMGL D3
Commercial iMGL D4
Gosselin Ex vivo
Olah Ex vivo
Galatro Ex vivo
Hasselmann iMGL
Svoboda iMGL
Hasselmann xMG
Svoboda xMG
Gosselin In vitro
Gosselin Monocyte

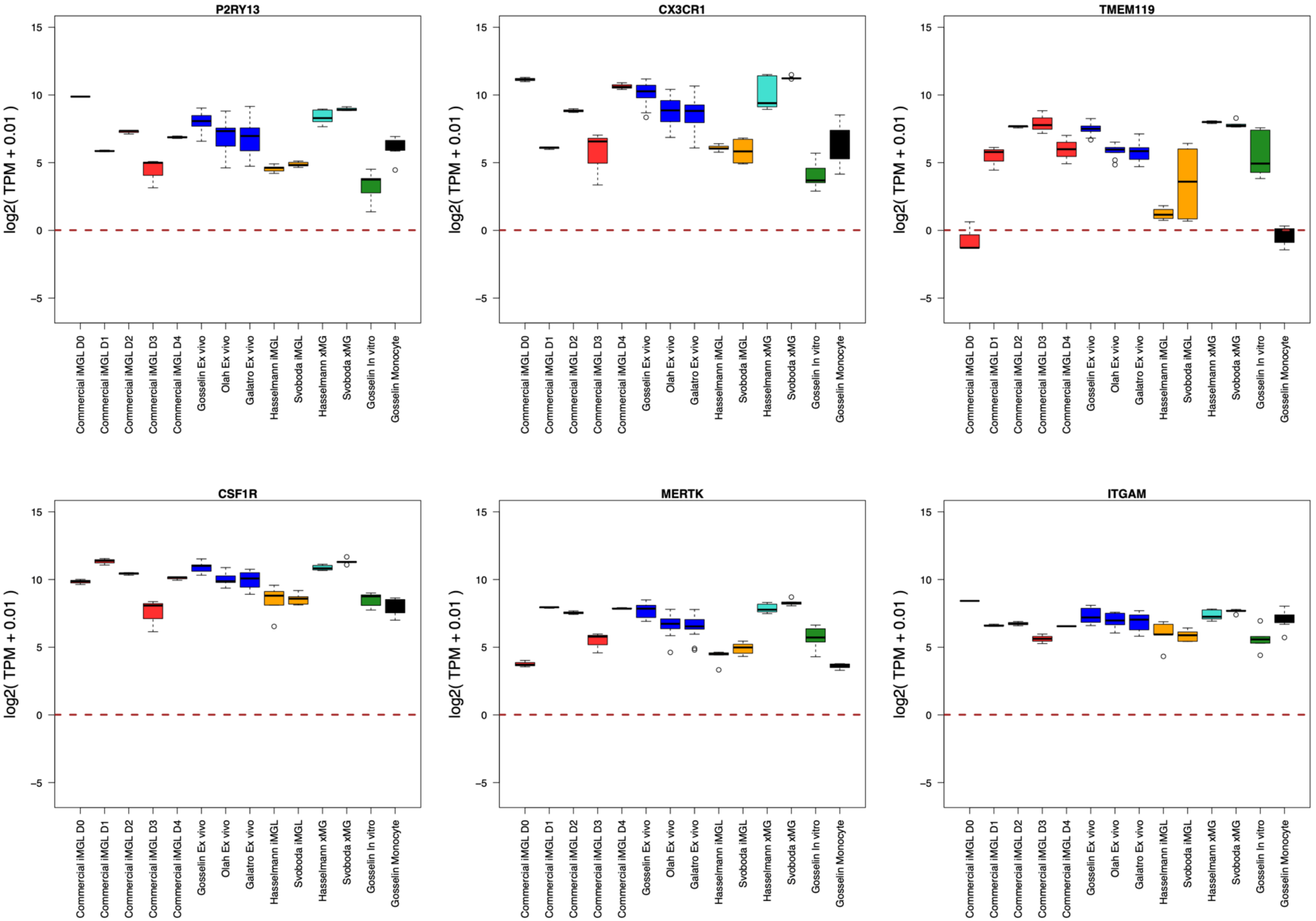

P2RY13
CX3CR1
TMEM119
CSF1R
MERTK
ITGAM
log2( TPM + 0.01 )
15
10
5
0
−5
Commercial iMGL D0
Commercial iMGL D1
Commercial iMGL D2
Commercial iMGL D3
Commercial iMGL D4
Gosselin Ex vivo
Olah Ex vivo
Galatro Ex vivo
Hasselmann iMGL
Svoboda iMGL
Hasselmann xMG
Svoboda xMG
Gosselin In vitro
Gosselin Monocyte

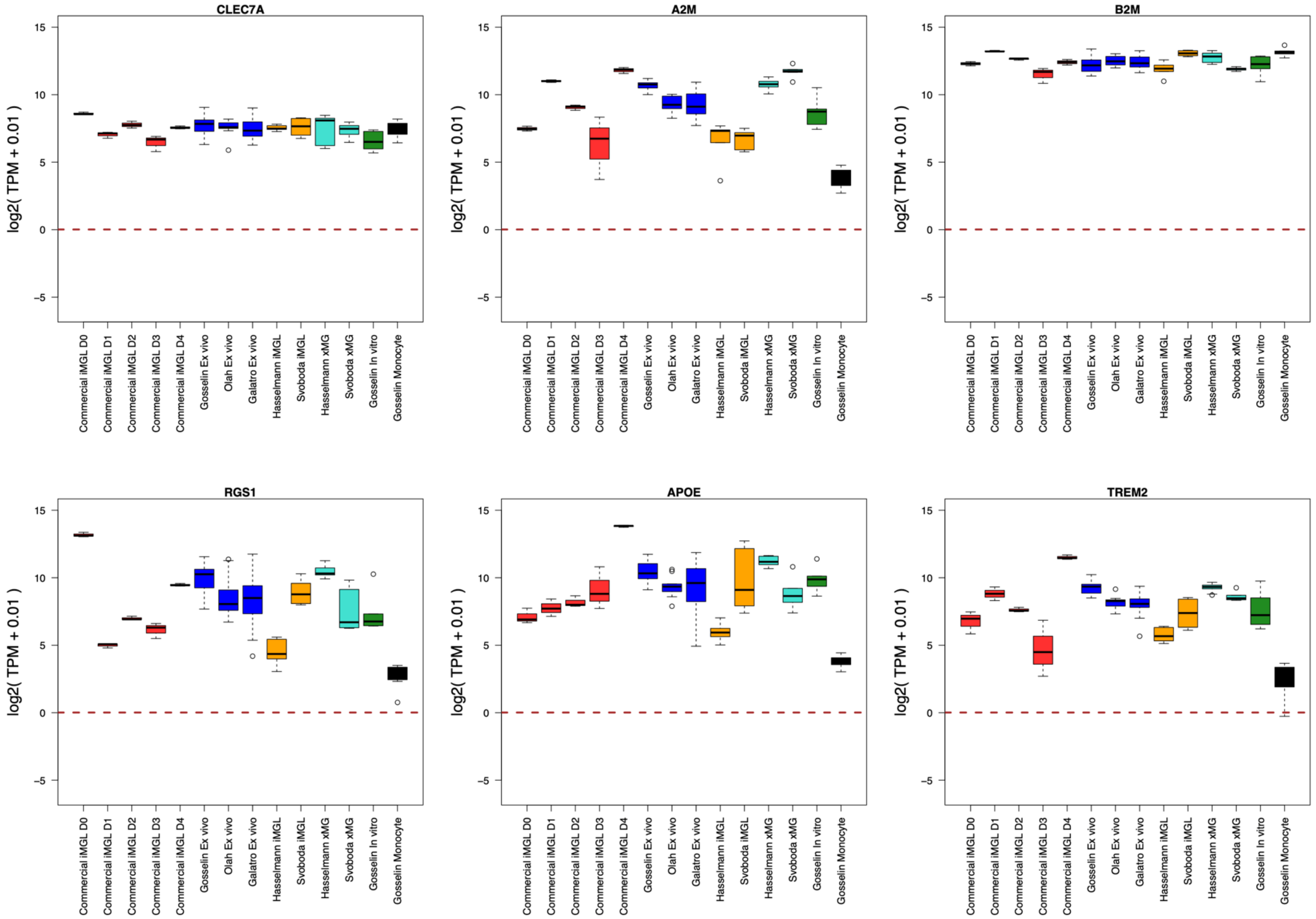
CLEC7A
A2M
B2M
RGS1
APOE
TREM2
log2( TPM + 0.01 )
Commercial iMGL D0
Commercial iMGL D1
Commercial iMGL D2
Commercial iMGL D3
Commercial iMGL D4
Gosselin Ex vivo
Olah Ex vivo
Galatro Ex vivo
Hasselmann iMGL
Svoboda iMGL
Hasselmann xMG
Svoboda xMG
Gosselin In vitro
Gosselin Monocyte

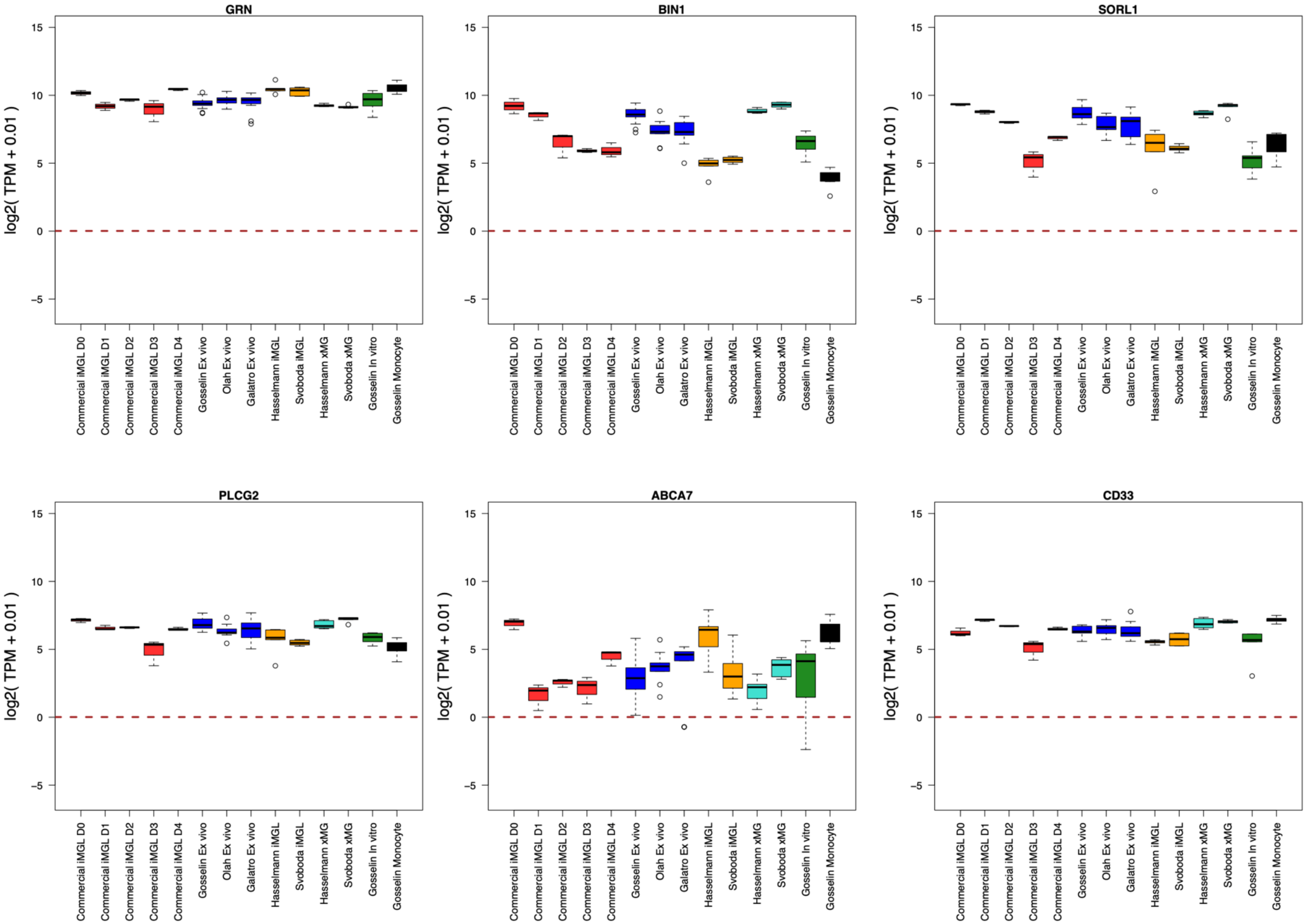
GRN
BIN1
SORL1
PLCG2
ABCA7
CD33
log2( TPM + 0.01 )
15
10
5
0
−5
Commercial iMGL D0
Commercial iMGL D1
Commercial iMGL D2
Commercial iMGL D3
Commercial iMGL D4
Gosselin Ex vivo
Olah Ex vivo
Galatro Ex vivo
Hasselmann iMGL
Svoboda iMGL
Hasselmann xMG
Svoboda xMG
Gosselin In vitro
Gosselin Monocyte

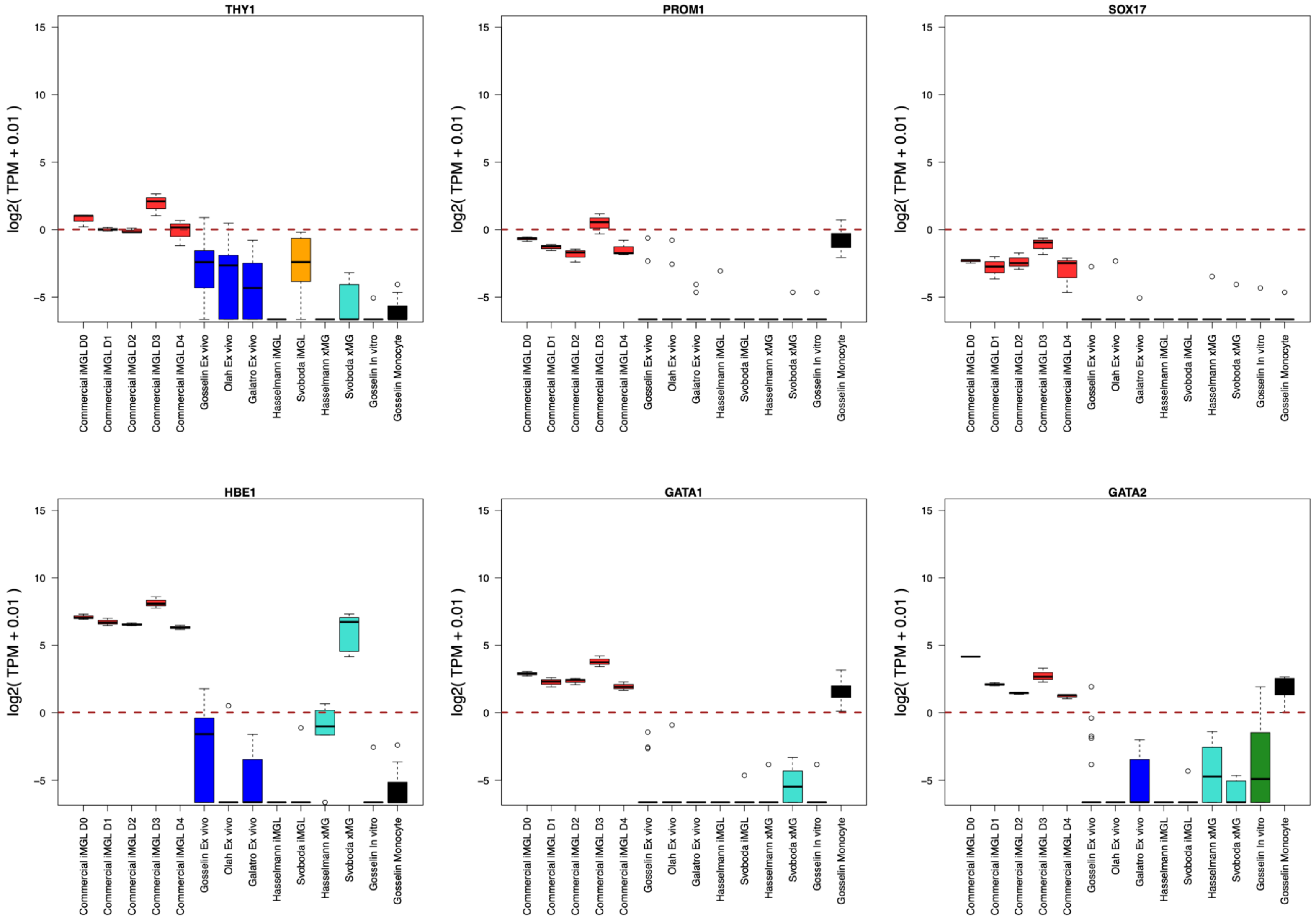
THY1
PROM1
SOX17
HBE1
GATA1
GATA2
log2( TPM + 0.01 )
15
10
5
0
-5
Commercial iMGL D0
Commercial iMGL D1
Commercial iMGL D2
Commercial iMGL D3
Commercial iMGL D4
Gosselin Ex vivo
Olah Ex vivo
Galatro Ex vivo
Hasselmann iMGL
Svoboda iMGL
Hasselmann xMG
Svoboda xMG
Gosselin In vitro
Gosselin Monocyte

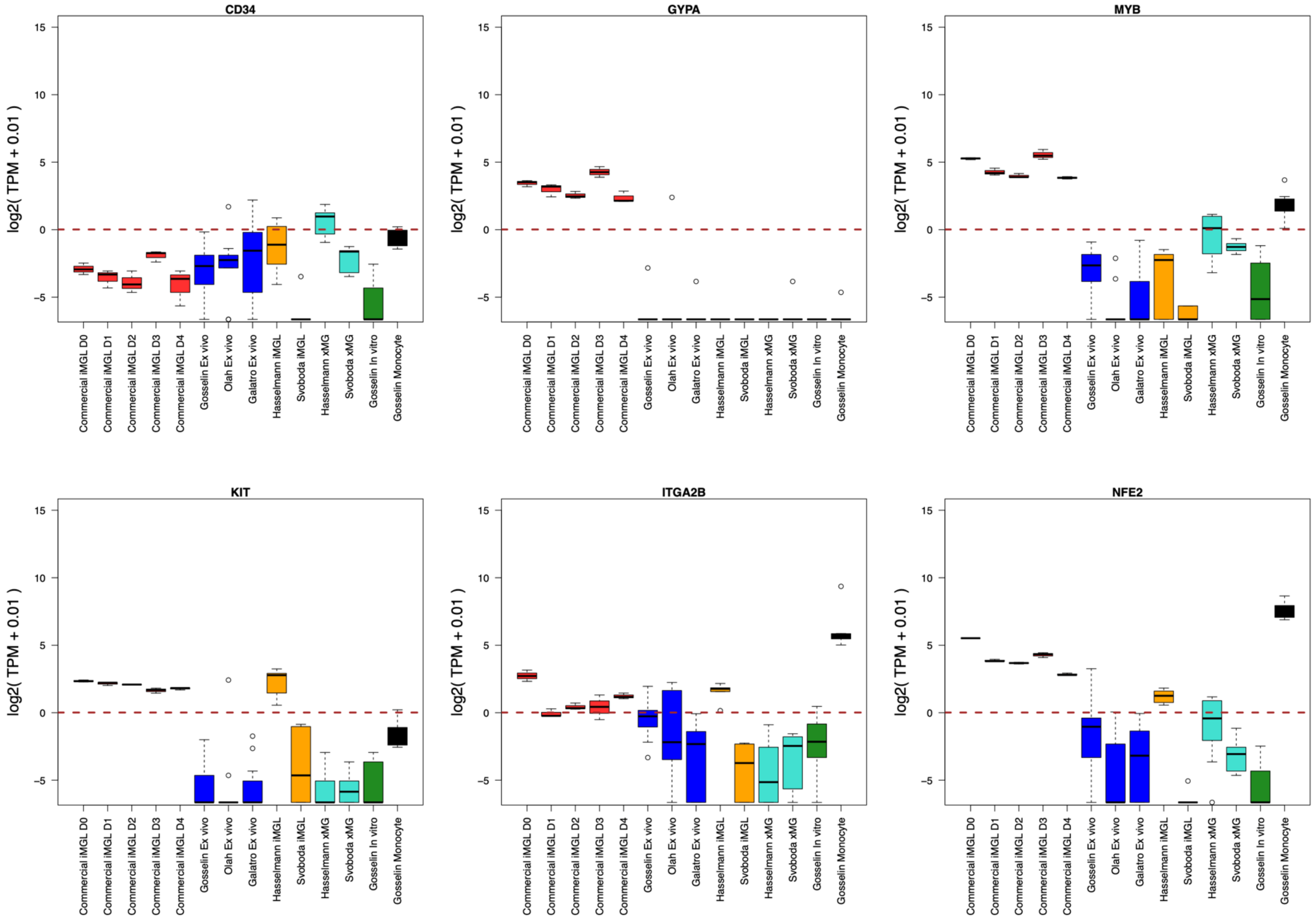

CD34
GYPA
MYB
KIT
ITGA2B
NFE2
log2( TPM + 0.01 )
15
10
5
0
−5
Commercial iMGL D0
Commercial iMGL D1
Commercial iMGL D2
Commercial iMGL D3
Commercial iMGL D4
Gosselin Ex vivo
Olah Ex vivo
Galatro Ex vivo
Hasselmann iMGL
Svoboda iMGL
Hasselmann xMG
Svoboda xMG
Gosselin In vitro
Gosselin Monocyte

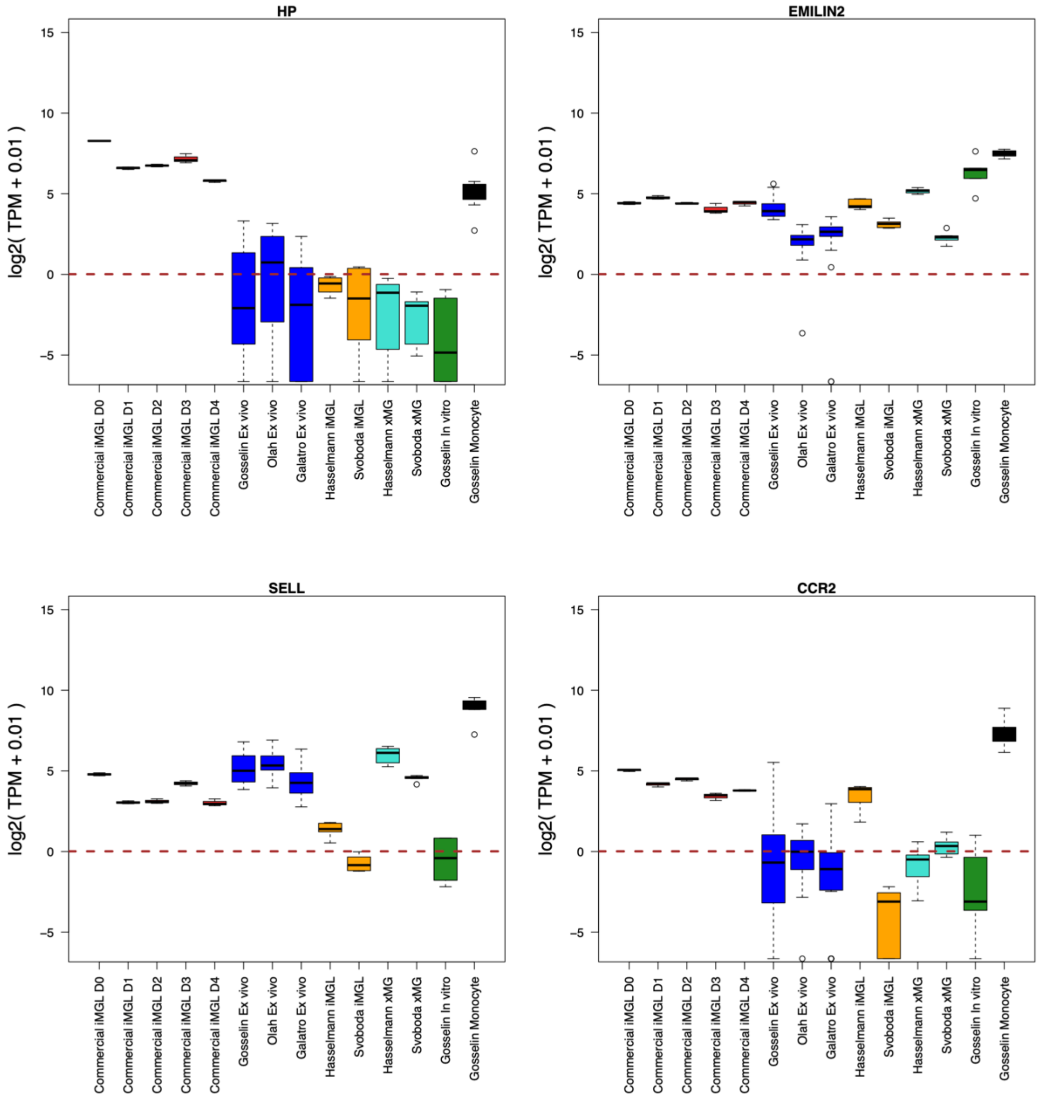

HP
EMILIN2
SELL
CCR2
log2( TPM + 0.01 )
Commercial iMGL D0
Commercial iMGL D1
Commercial iMGL D2
Commercial iMGL D3
Commercial iMGL D4
Gosselin Ex vivo
Olah Ex vivo
Galatro Ex vivo
Hasselmann iMGL
Svoboda iMGL
Hasselmann xMG
Svoboda xMG
Gosselin In vitro
Gosselin Monocyte

**Supplementary Figure 3.** Expression of key microglia related genes across the Commercial iMGL samples and microglia comparator datasets.

Microglia TFs: SPI1, IRF8, RUNX1, FOS, CEBPB, MEF2A, MAF, SALL1, TAL1, RUNX2

Microglia core marker genes: C3, C1QA, C1QB, C1QC, HLA-B. HLA-DRA, SPP1, P2RY12, P2RY13, CX3CR1, TMEM119, CSF1R, MERTK, ITGAM, CLEC7A, A2M, B2M, RGS1

Disease genes: APOE, TREM2, GRN, BIN1, SORL1, PLCG2, ABCA7, CD33

Myeloid progenitor and monocyte marker genes: THY1, PROM1, SOX17, HBE1, GATA1, GATA2, CD34, GYPA, MYB, KIT, ITGA2B, NFE2, HP, EMILIN2, SELL, CCR2

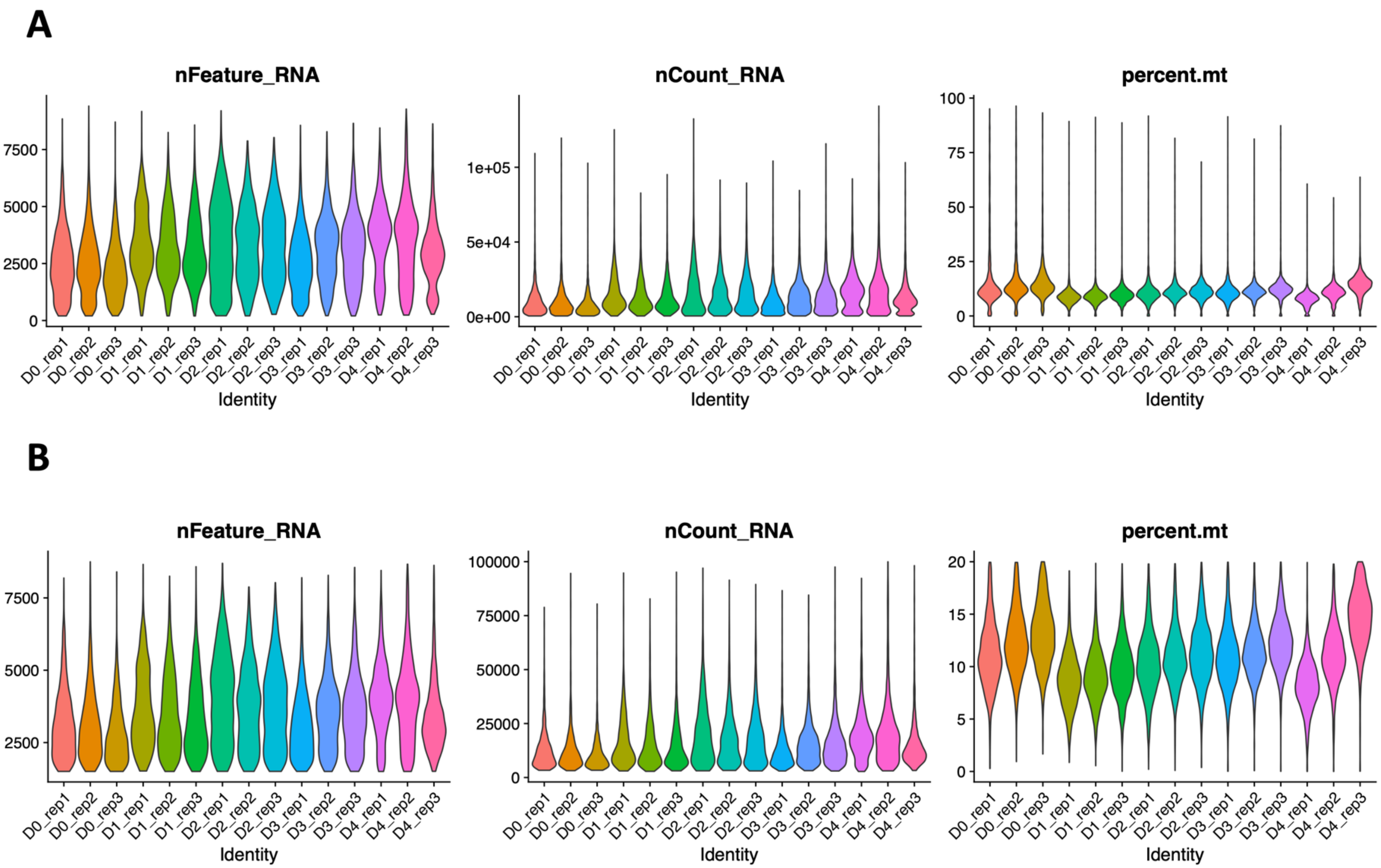
A
nFeature_RNA
nCount_RNA
percent.mt
B
nFeature_RNA
nCount_RNA
percent.mt
Identity
D0_rep1
D0_rep2
D0_rep3
D1_rep1
D1_rep2
D1_rep3
D2_rep1
D2_rep2
D2_rep3
D3_rep1
D3_rep2
D3_rep3
D4_rep1
D4_rep2
D4_rep3

**Supplementary Figure 4.** Quality control (QC) metrics for Commercial iMGL time course single cell RNA-seq dataset. The number of genes (nFeature_RNA), reads (nCount_RNA), and percentage of mitochondrial reads (percent.mt) for cells in each sample **A**) before and **B**) after applying QC filters.

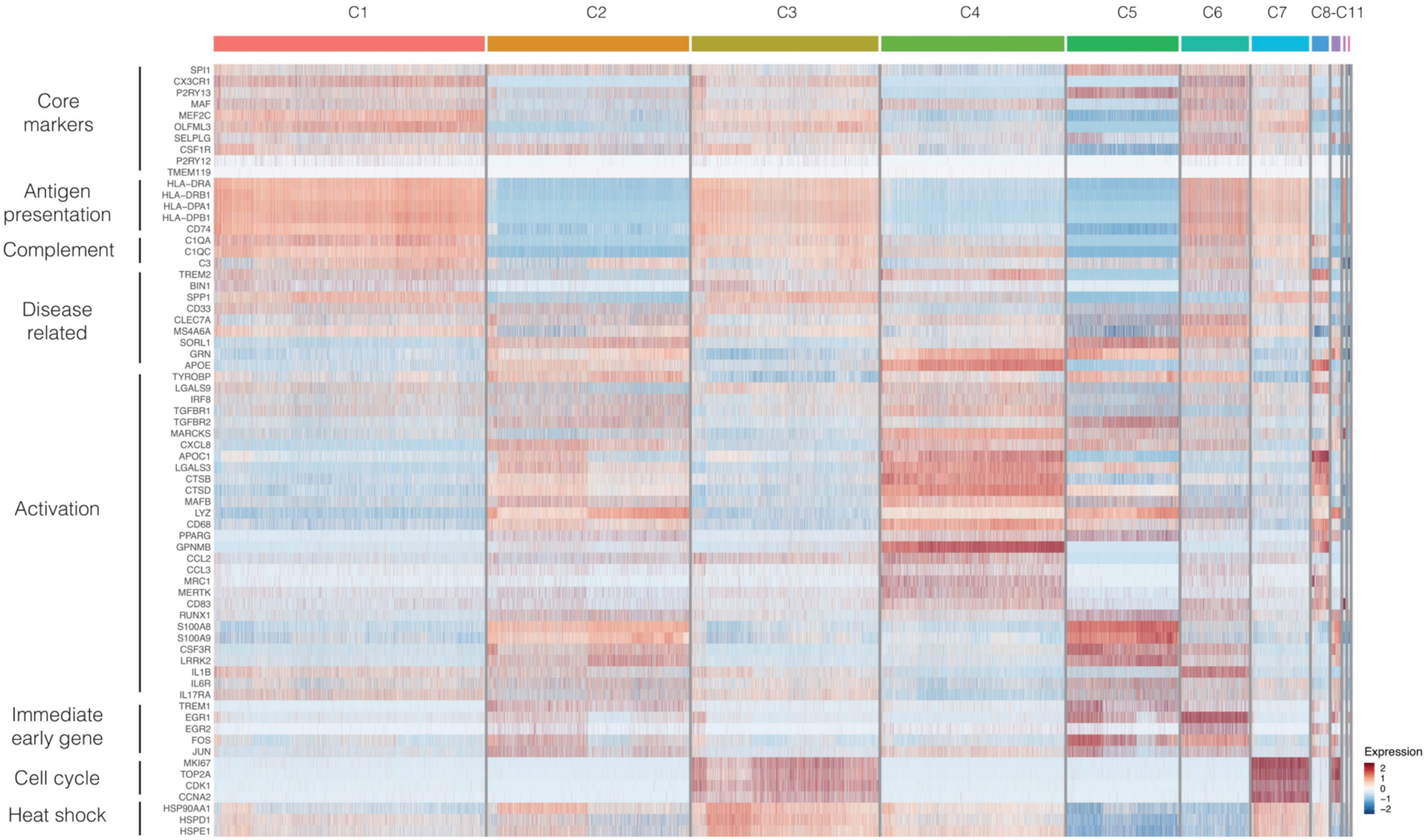
C1
C2
C3
C4
C5
C6
C7
C8-C11
Core markers
Antigen presentation
Complement
Disease related
Activation
Immediate early gene
Cell cycle
Heat shock
SPI1
CX3CR1
P2RY13
MAF
MEF2C
OLFML3
SELPLG
CSF1R
P2RY12
TMEM119
HLA-DRA
HLA-DRB1
HLA-DPA1
HLA-DPB1
CD74
C1QA
C1QC
C3
TREM2
BIN1
SPP1
CD33
CLEC7A
MS4A6A
SORL1
GRN
APOE
TYROBP
LGALS9
IRF8
TGFBR1
TGFBR2
MARCKS
CXCL8
APOC1
LGALS3
CTSB
CTSD
MAFB
LYZ
CD68
PPARG
GPNMB
CCL2
CCL3
MRC1
MERTK
CD83
RUNX1
S100A8
S100A9
CSF3R
LRRK2
IL1B
IL6R
IL17RA
TREM1
EGR1
EGR2
FOS
JUN
MKI67
TOP2A
CDK1
CCNA2
HSP90AA1
HSPD1
HSPE1
Expression
2
1
0
−1
−2

**Supplementary Figure 5.** Gene expression heatmap of microglia functional marker genes in the time course single cell RNA-seq dataset.

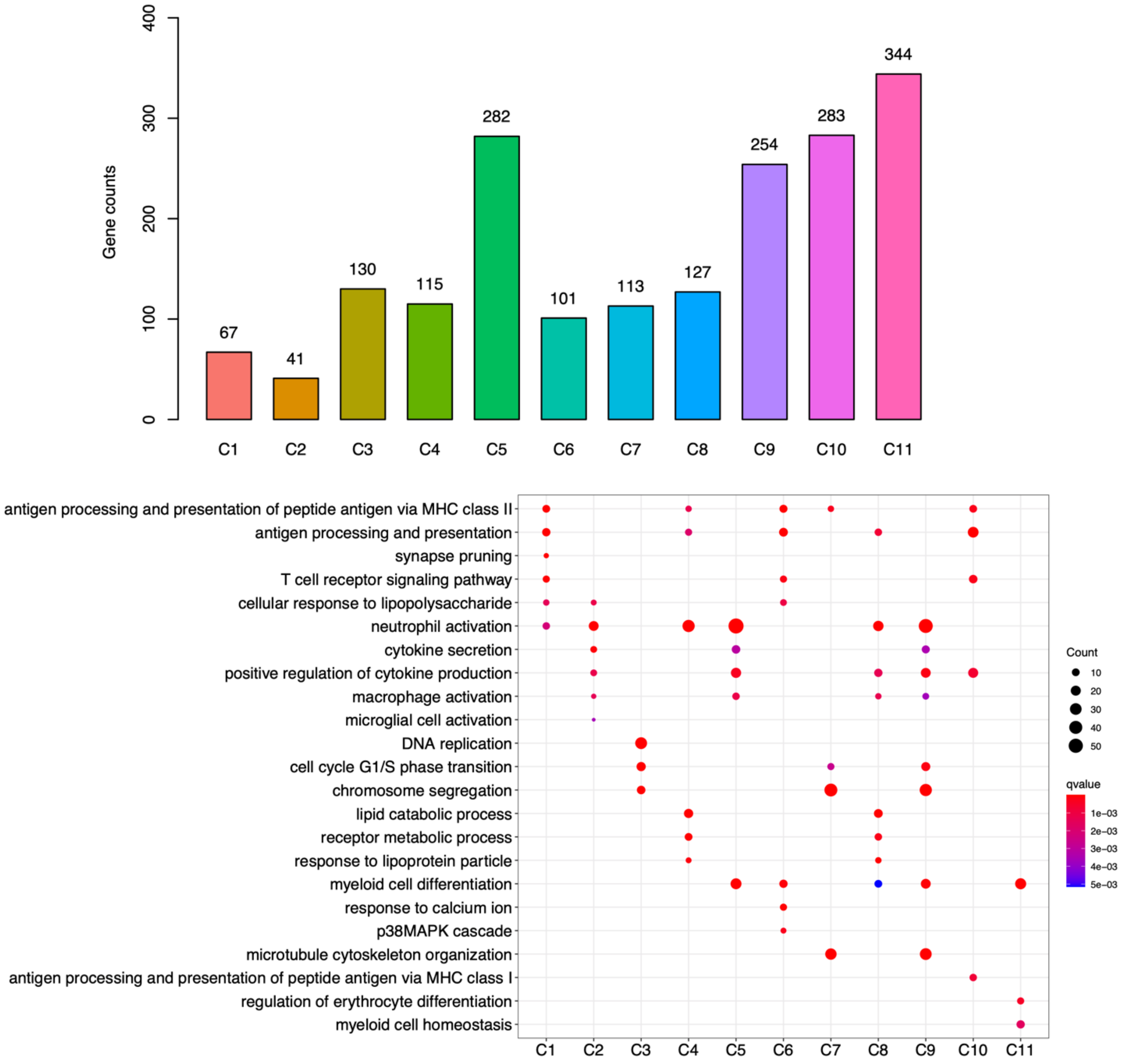

Gene counts
0
100
200
300
400
67
41
130
115
282
101
113
127
254
283
344
C1
C2
C3
C4
C5
C6
C7
C8
C9
C10
C11
antigen processing and presentation of peptide antigen via MHC class II
antigen processing and presentation
synapse pruning
T cell receptor signaling pathway
cellular response to lipopolysaccharide
neutrophil activation
cytokine secretion
positive regulation of cytokine production
macrophage activation
microglial cell activation
DNA replication
cell cycle G1/S phase transition
chromosome segregation
lipid catabolic process
receptor metabolic process
response to lipoprotein particle
myeloid cell differentiation
response to calcium ion
p38MAPK cascade
microtubule cytoskeleton organization
antigen processing and presentation of peptide antigen via MHC class I
regulation of erythrocyte differentiation
myeloid cell homeostasis
Count
10
20
30
40
50
qvalue
1e−03
2e−03
3e−03
4e−03
5e−03

**Supplementary Figure 6.** Identification of preferentially expressed marker genes and their top gene ontology enrichments for each cellular cluster in the time course single cell RNA-seq dataset.

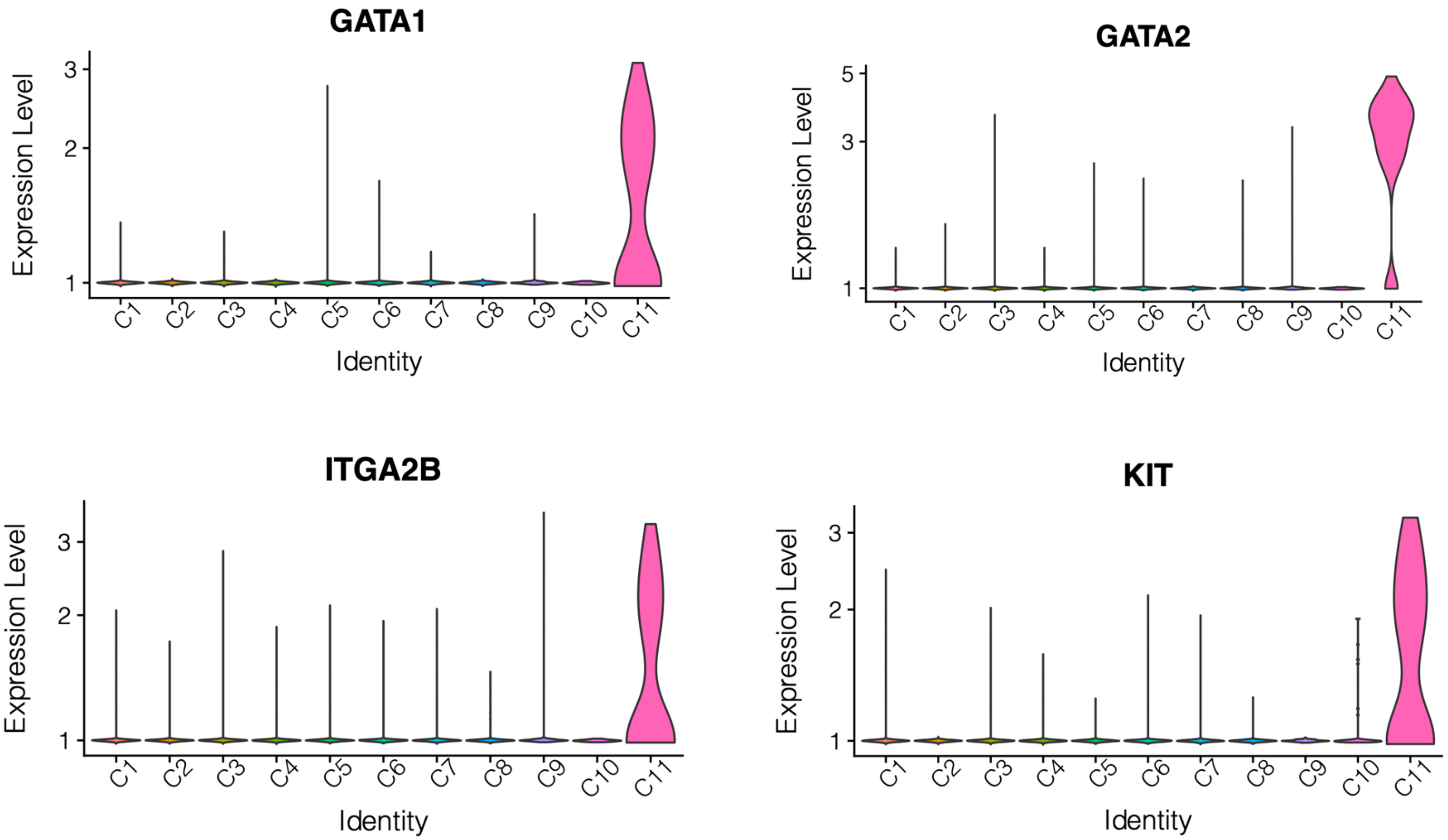
GATA1
GATA2
ITGA2B
KIT
Expression Level
Identity
C1 C2 C3 C4 C5 C6 C7 C8 C9 C10 C11

**Supplementary Figure 7.** Expression of progenitor (GATA1, GATA2, KIT) and monocyte (ITGA2B) marker genes across the 11 single cell clusters.

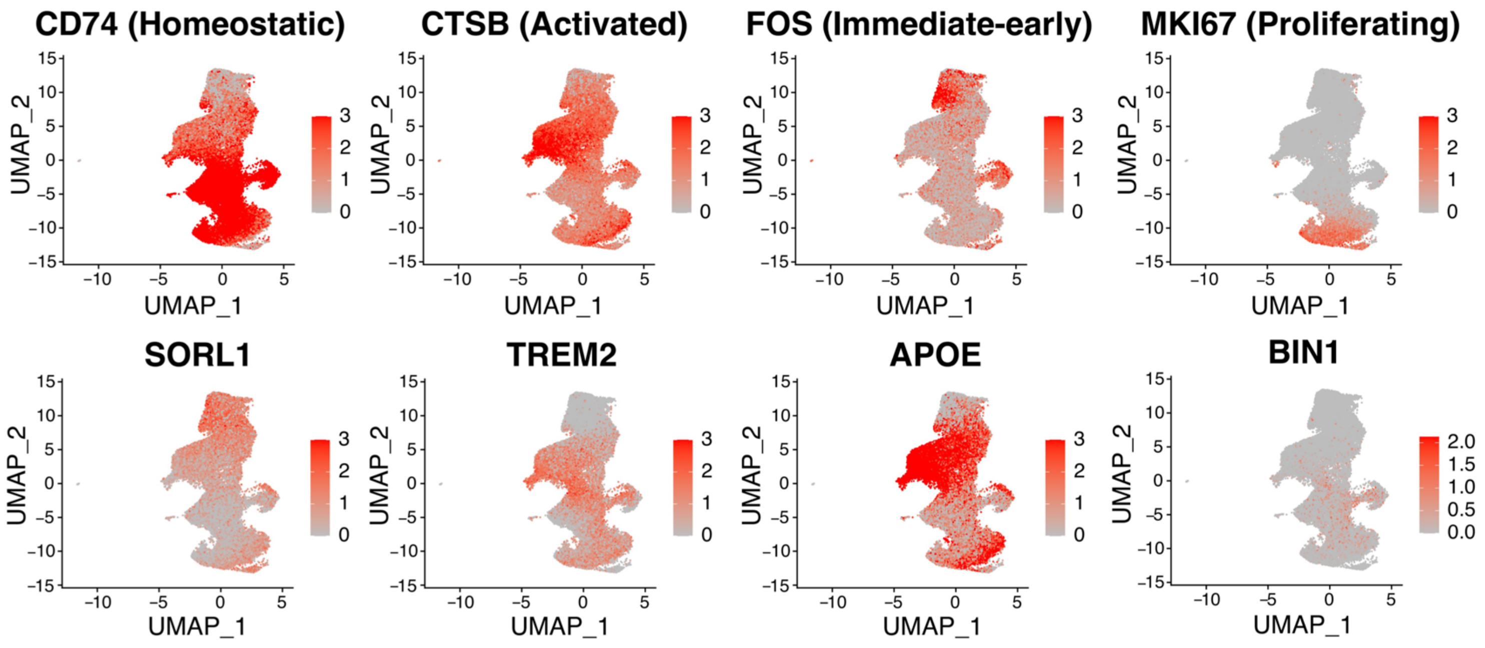
CD74 (Homeostatic)
CTSB (Activated)
FOS (Immediate-early)
MKI67 (Proliferating)
SORL1
TREM2
APOE
BIN1
UMAP_1
UMAP_2

**Supplementary Figure 8.** Relative expression of key homeostatic (CD74), activated (CTSB), immediate-early (FOS), proliferating (MKI67), and AD-associated (SORL1, TREM2, APOE, BIN1) genes across the individual cells. Expression values were scaled using the FeaturePlot method in Seurat.

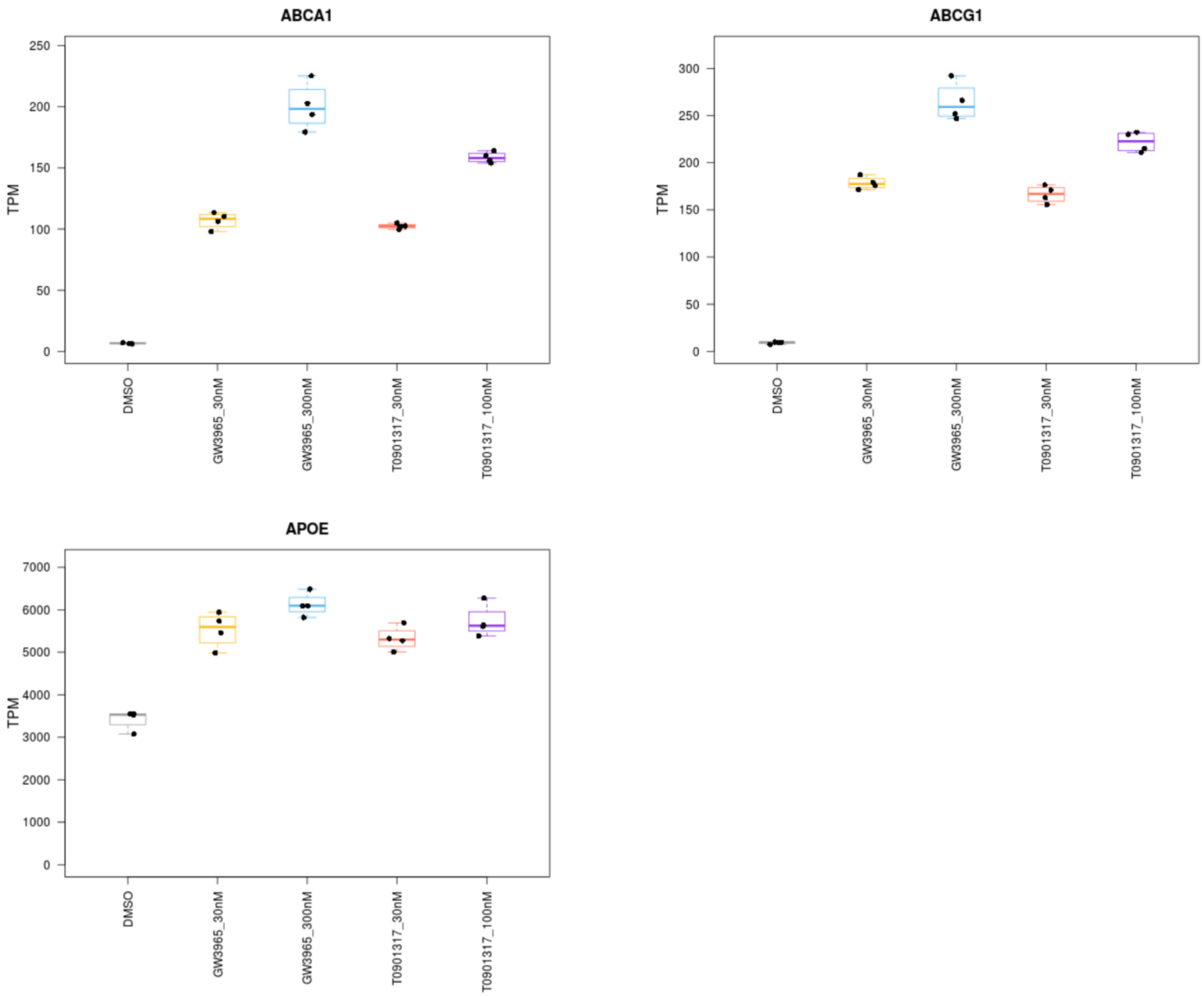
ABCA1
ABCG1
APOE
TPM
DMSO
GW3965_30nM
GW3965_300nM
T0901317_30nM
T0901317_100nM
0
50
100
150
200
250
300
1000
2000
3000
4000
5000
6000
7000

**Supplementary Figure 9.** Expression of known LXR pathway response genes (ABCA1, ABCG1, APOE) in the LXR pathway agonist treatment bulk RNA-seq dataset.

GW3965 30nM:

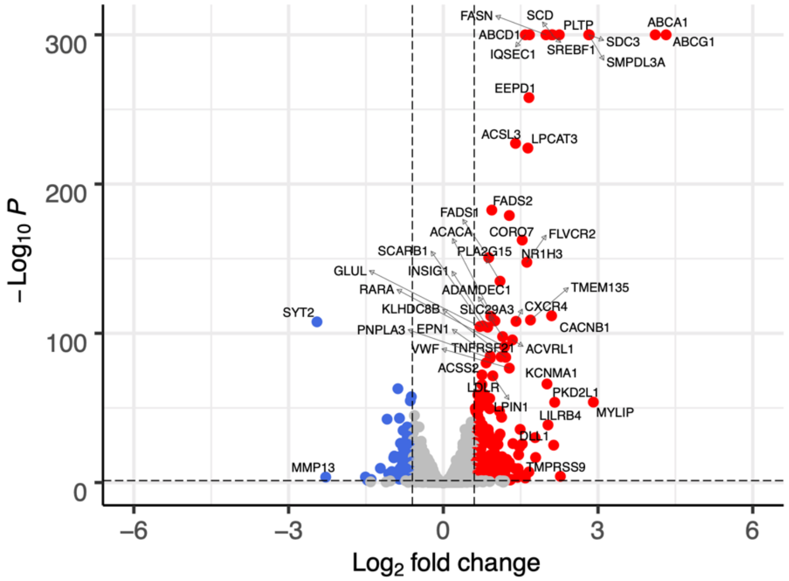


T0901317 30nM:

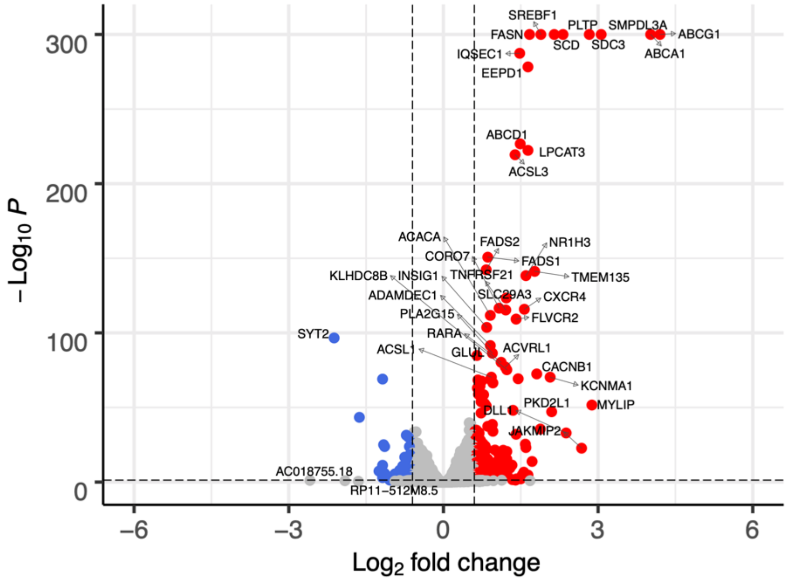

**Supplementary Figure 10.** Volcano plots showing relationship between expression fold change and significance for differential gene expression between GW3965 30nM and T0901317 30nM with DMSO treated cells. Genes with significant upregulation and downregulation are colored red and blue, respectively.

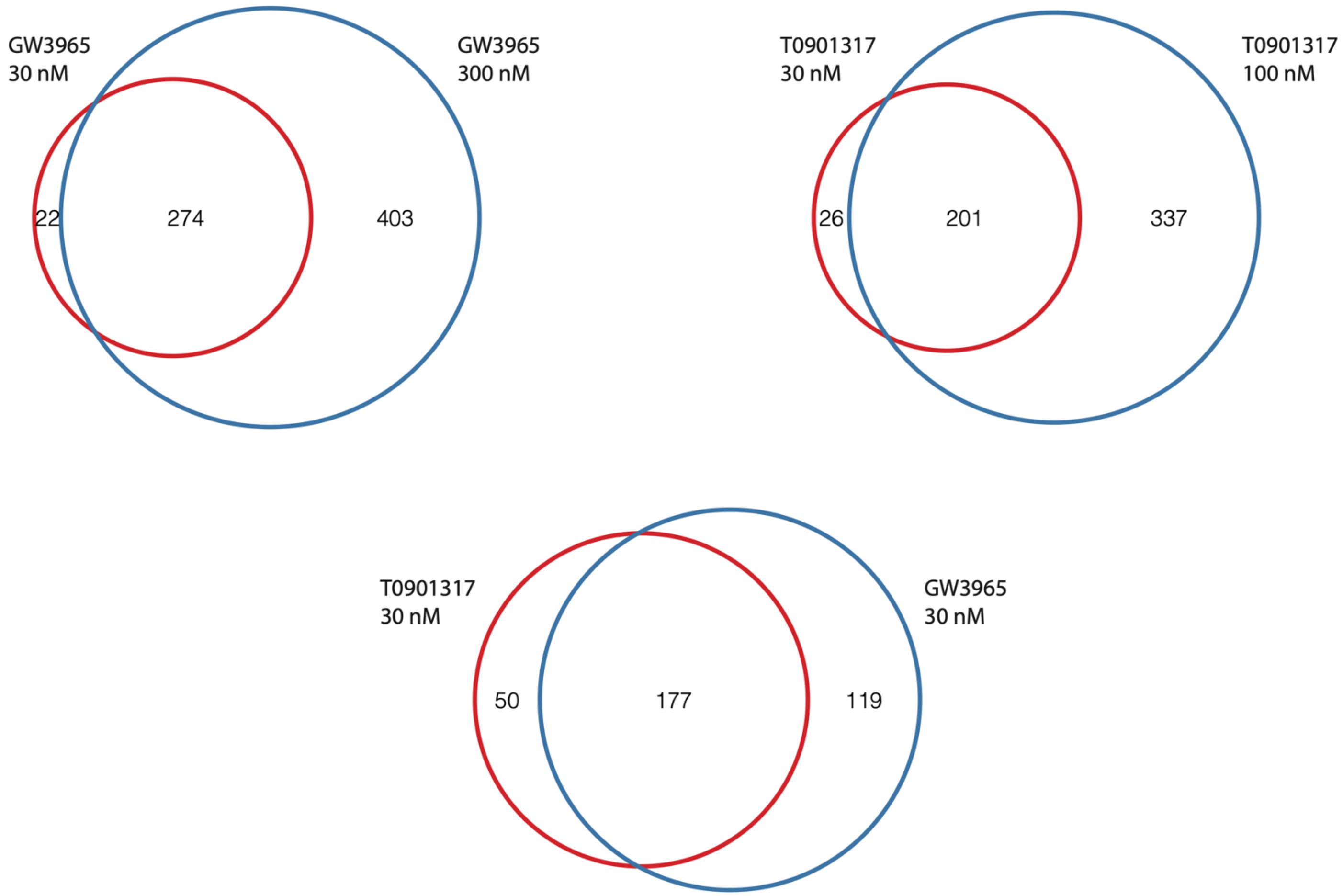
GW3965
30 nM
GW3965
300 nM
22
274
403
T0901317
30 nM
T0901317
100 nM
26
201
337
T0901317
30 nM
GW3965
30 nM
50
177
119

**Supplementary Figure 11.** Overlap in DEGs between GW3965 30nM and 300nM treatments. Overlap in DEGs between T0901317 30nM and 100nM treatments. Overlap in DEGs between GW3965 30nM and T0901317 30nM treatments.

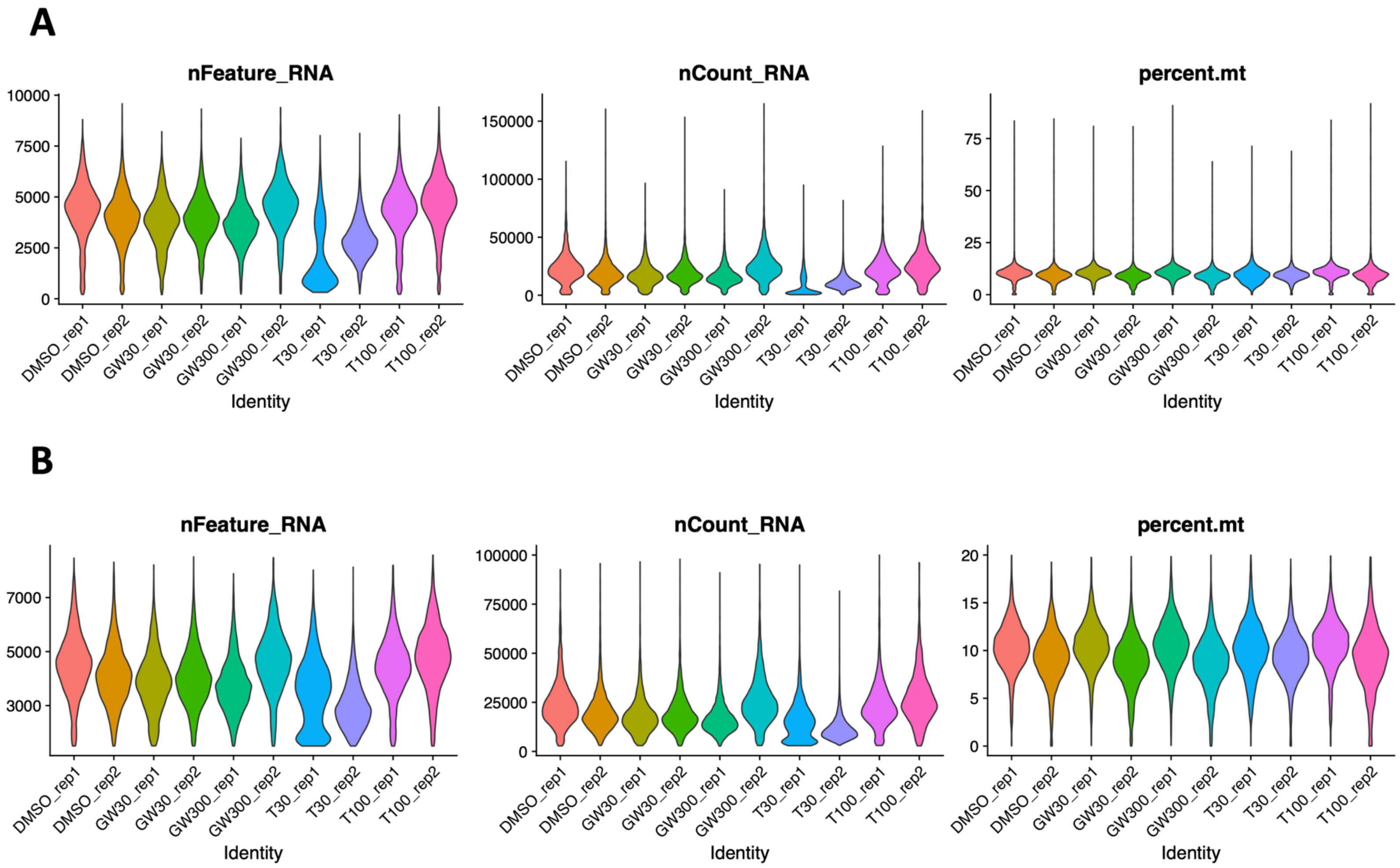
A
nFeature_RNA
nCount_RNA
percent.mt
10000
7500
5000
2500
0
150000
100000
50000
0
75
50
25
0
DMSO_rep1
DMSO_rep2
GW30_rep1
GW30_rep2
GW300_rep1
GW300_rep2
T30_rep1
T30_rep2
T100_rep1
T100_rep2
Identity
B
nFeature_RNA
nCount_RNA
percent.mt
7000
5000
3000
100000
75000
50000
25000
0
20
15
10
5
0
DMSO_rep1
DMSO_rep2
GW30_rep1
GW30_rep2
GW300_rep1
GW300_rep2
T30_rep1
T30_rep2
T100_rep1
T100_rep2
Identity

**Supplementary Figure 12.** QC metrics for LXR pathway agonist treatment single cell RNA-seq dataset. The number of genes (nFeature_RNA), reads (nCount_RNA), and percentage of mitochondrial reads (percent.mt) for cells for cells in each sample **A**) before and **B**) after applying QC filters.

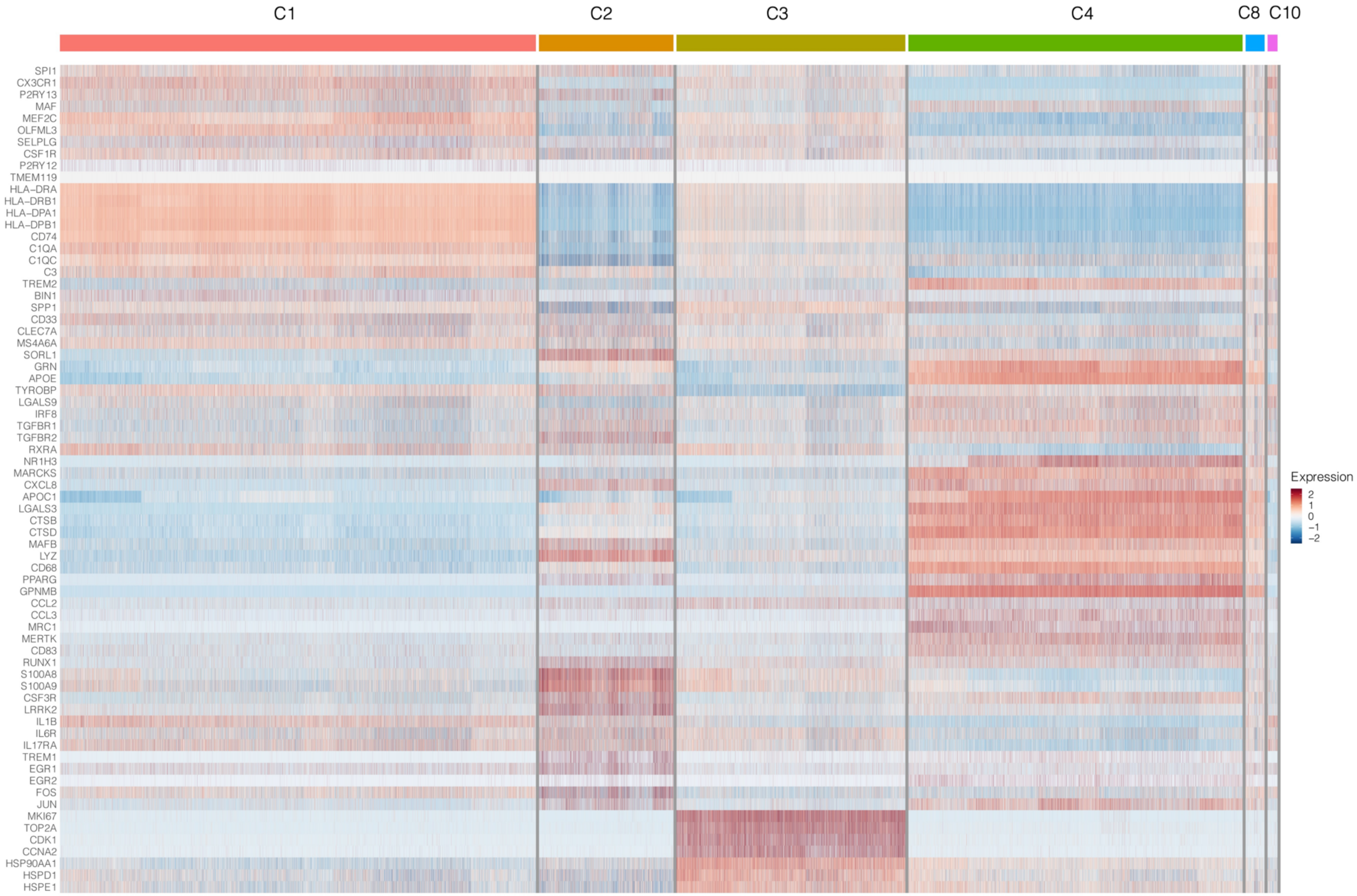
C1
C2
C3
C4
C8
C10
SPI1
CX3CR1
P2RY13
MAF
MEF2C
OLFML3
SELPLG
CSF1R
P2RY12
TMEM119
HLA−DRA
HLA−DRB1
HLA−DPA1
HLA−DPB1
CD74
C1QA
C1QC
C3
TREM2
BIN1
SPP1
CD33
CLEC7A
MS4A6A
SORL1
GRN
APOE
TYROBP
LGALS9
IRF8
TGFBR1
TGFBR2
RXRA
NR1H3
MARCKS
CXCL8
APOC1
LGALS3
CTSB
CTSD
MAFB
LYZ
CD68
PPARG
GPNMB
CCL2
CCL3
MRC1
MERTK
CD83
RUNX1
S100A8
S100A9
CSF3R
LRRK2
IL1B
IL6R
IL17RA
TREM1
EGR1
EGR2
FOS
JUN
MKI67
TOP2A
CDK1
CCNA2
HSP90AA1
HSPD1
HSPE1
Expression
2
1
0
−1
−2

**Supplementary Figure 13.** Gene expression heatmap of microglia functional marker genes in the LXR pathway agonist treatment single cell RNA-seq dataset.

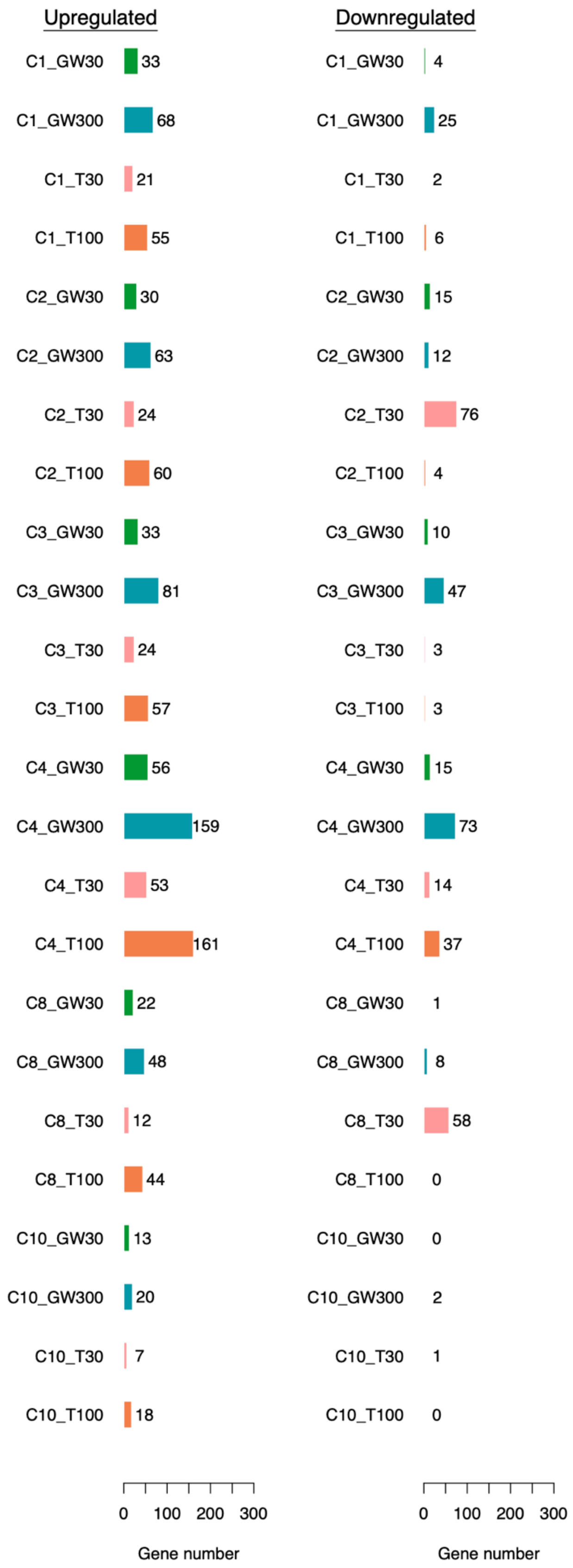

Upregulated
Downregulated
C1_GW30 33
C1_GW300 68
C1_T30 21
C1_T100 55
C2_GW30 30
C2_GW300 63
C2_T30 24
C2_T100 60
C3_GW30 33
C3_GW300 81
C3_T30 24
C3_T100 57
C4_GW30 56
C4_GW300 159
C4_T30 53
C4_T100 161
C8_GW30 22
C8_GW300 48
C8_T30 12
C8_T100 44
C10_GW30 13
C10_GW300 20
C10_T30 7
C10_T100 18
C1_GW30 4
C1_GW300 25
C1_T30 2
C1_T100 6
C2_GW30 15
C2_GW300 12
C2_T30 76
C2_T100 4
C3_GW30 10
C3_GW300 47
C3_T30 3
C3_T100 3
C4_GW30 15
C4_GW300 73
C4_T30 14
C4_T100 37
C8_GW30 1
C8_GW300 8
C8_T30 58
C8_T100 0
C10_GW30 0
C10_GW300 2
C10_T30 1
C10_T100 0
0 100 200 300
Gene number
0 100 200 300
Gene number

**Supplementary Figure 14.** Number of DEGs identified in each single cell cluster for each treatment condition against DMSO. GW30: GW3965 30 nM, GW300: GW3965 300 nM, T30: T0901317 30 nM, T100: T0901317 100 nM.

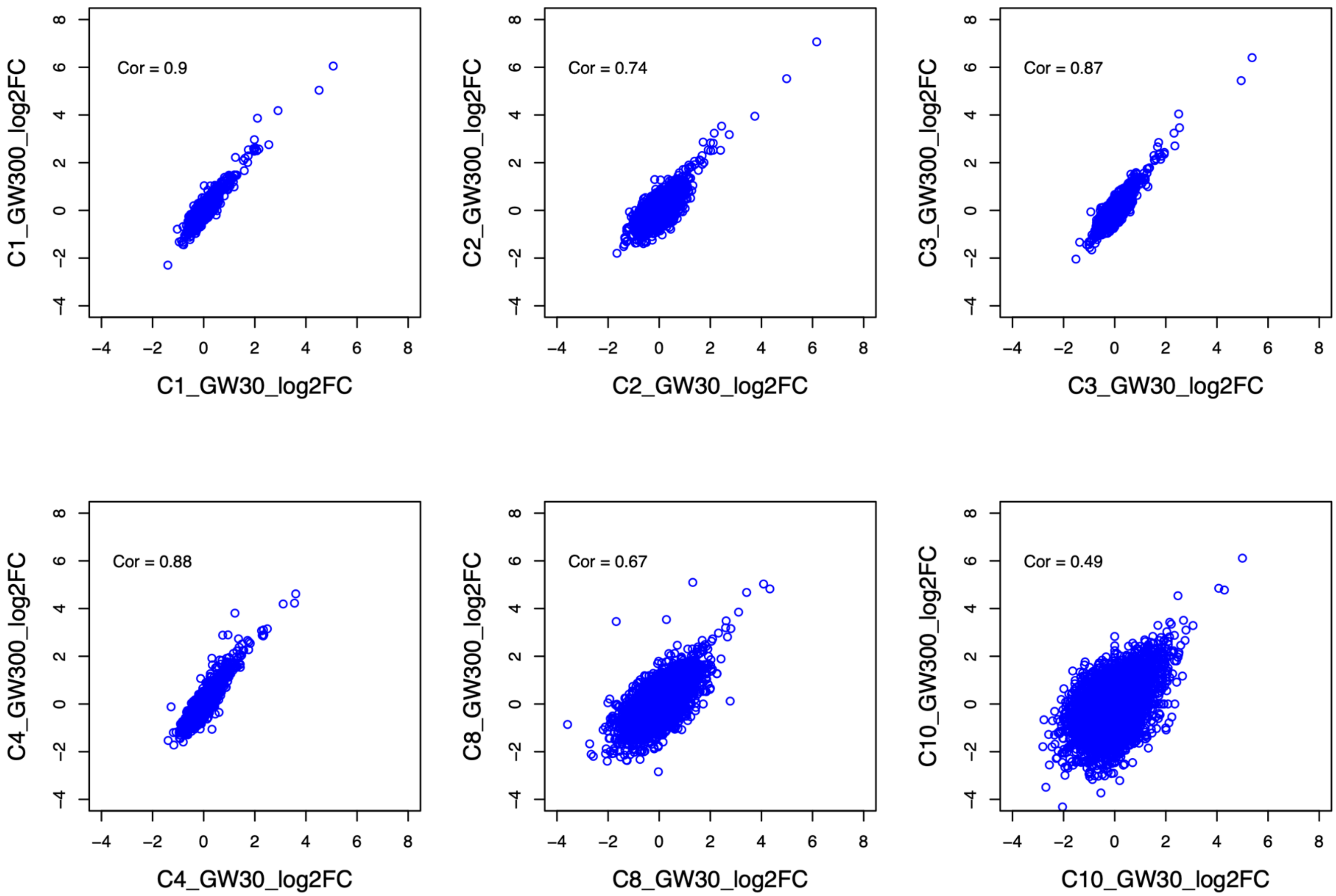
Cor = 0.9
Cor = 0.74
Cor = 0.87
Cor = 0.88
Cor = 0.67
Cor = 0.49
C1_GW300_log2FC
C1_GW30_log2FC
C2_GW300_log2FC
C2_GW30_log2FC
C3_GW300_log2FC
C3_GW30_log2FC
C4_GW300_log2FC
C4_GW30_log2FC
C8_GW300_log2FC
C8_GW30_log2FC
C10_GW300_log2FC
C10_GW30_log2FC

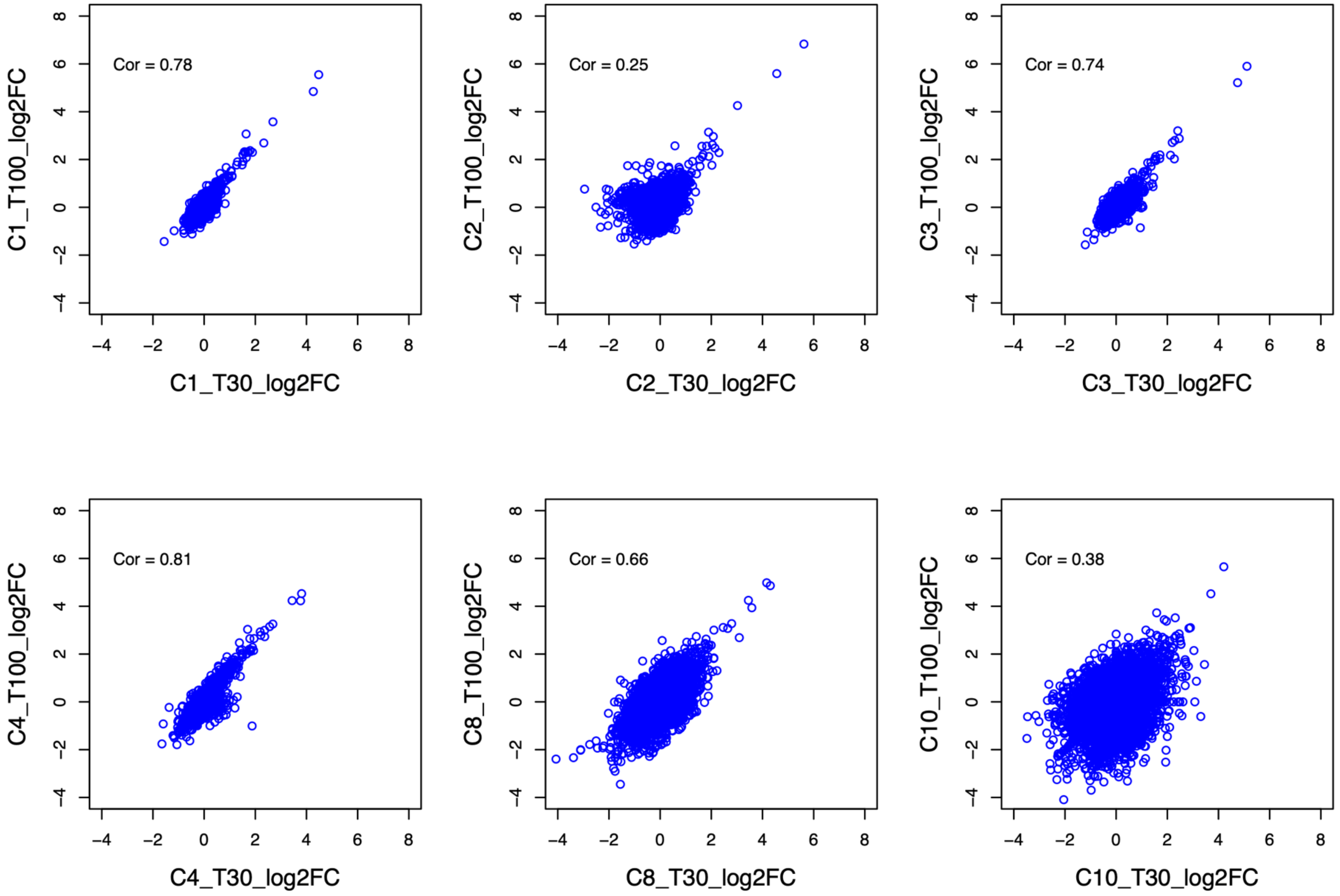
Cor = 0.78
C1_T30_log2FC
C1_T100_log2FC
Cor = 0.25
C2_T30_log2FC
C2_T100_log2FC
Cor = 0.74
C3_T30_log2FC
C3_T100_log2FC
Cor = 0.81
C4_T30_log2FC
C4_T100_log2FC
Cor = 0.66
C8_T30_log2FC
C8_T100_log2FC
Cor = 0.38
C10_T30_log2FC
C10_T100_log2FC

**Supplementary Figure 15.** Correlation of gene fold changes between the low and high dose of each treatment within the same single cell cluster. Pearson's correlations are shown. GW30: GW3965 30 nM, GW300: GW3965 300 nM, T30: T0901317 30 nM, T100: T0901317 100 nM.

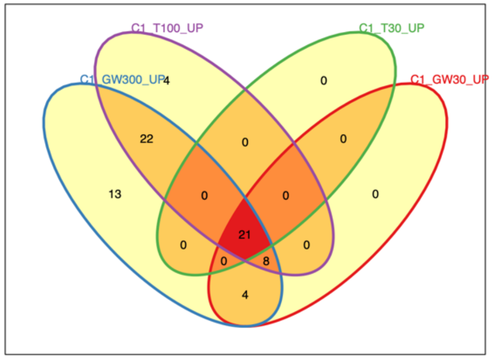
C1_T100_UP
C1_T30_UP
C1_GW300_UP
C1_GW30_UP
4
0
22
0
0
13
0
0
0
21
0
0
8
0
4

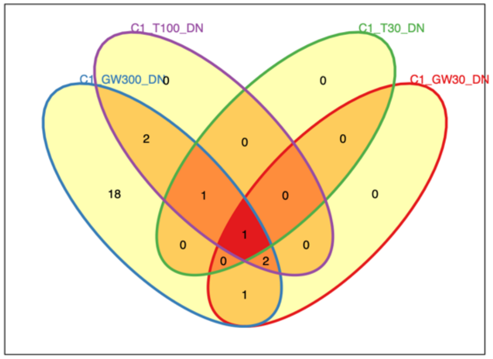
C1_T100_DN
C1_T30_DN
C1_GW300_DN
C1_GW30_DN
0
0
2
0
0
18
1
0
0
1
0
0
2
0
1

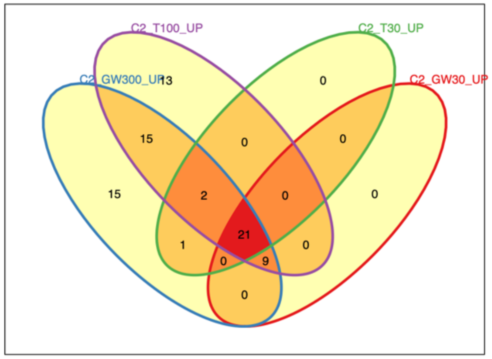
C2_T100_UP
C2_T30_UP
C2_GW300_UP
C2_GW30_UP
13
0
15
0
0
15
2
0
0
21
1
0
9
0
0

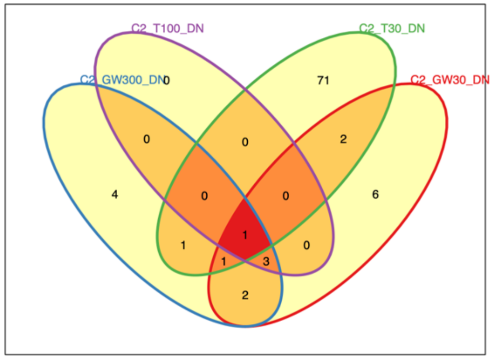
C2_T100_DN
C2_T30_DN
C2_GW300_DN
C2_GW30_DN
0
71
0
0
2
4
0
0
6
1
1
1
3
0
2

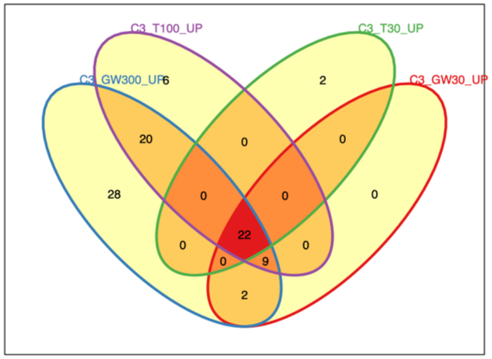
C3_T100_UP
C3_T30_UP
C3_GW300_UP
C3_GW30_UP
6
2
20
0
0
28
0
0
0
22
0
0
9
0
2

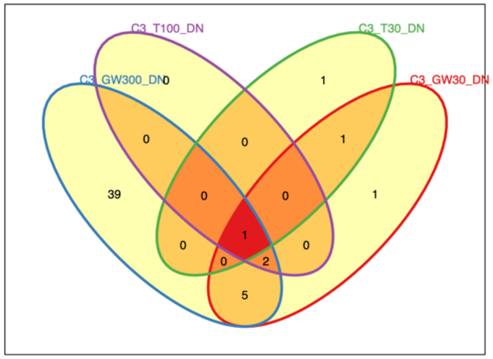
C3_T100_DN
C3_T30_DN
C3_GW300_DN
C3_GW30_DN
0
1
0
0
1
39
0
0
1
1
0
0
2
0
5

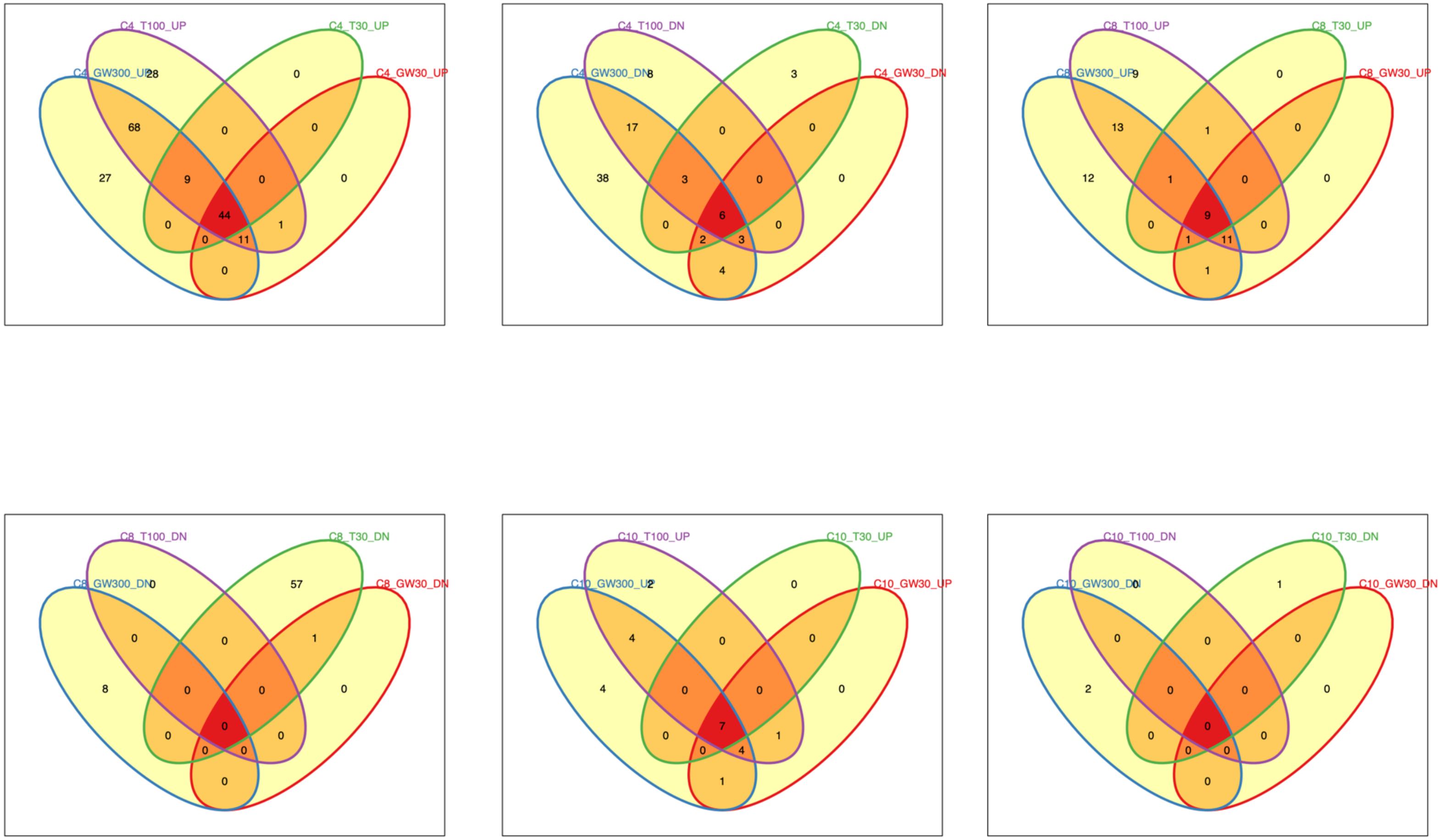
C4_T100_UP
C4_T30_UP
C4_GW300_UP
C4_GW30_UP
C4_T100_DN
C4_T30_DN
C4_GW300_DN
C4_GW30_DN
C8_T100_UP
C8_T30_UP
C8_GW300_UP
C8_GW30_UP
C8_T100_DN
C8_T30_DN
C8_GW300_DN
C8_GW30_DN
C10_T100_UP
C10_T30_UP
C10_GW300_UP
C10_GW30_UP
C10_T100_DN
C10_T30_DN
C10_GW300_DN
C10_GW30_DN

**Supplementary Figure 16.** Overlap in DEGs identified in each treatment within the same single cell cluster. UP: upregulated in the treatment compared to DMSO, DN: downregulated in the treatment compared to DMSO. GW30: GW3965 30 nM, GW300: GW3965 300 nM, T30: T0901317 30 nM, T100: T0901317 100 nM.

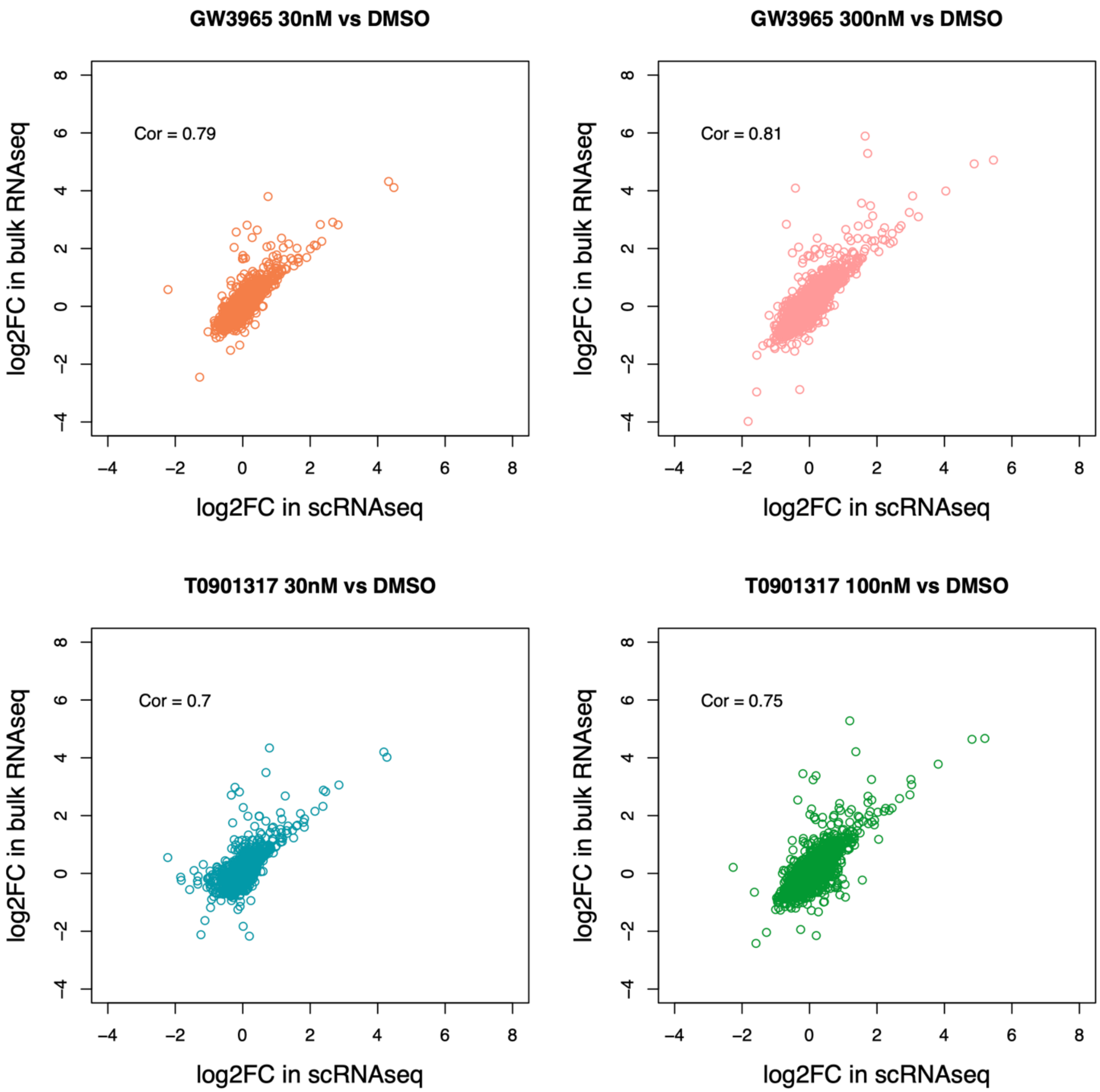
GW3965 30nM vs DMSO
Cor = 0.79
GW3965 300nM vs DMSO
Cor = 0.81
T0901317 30nM vs DMSO
Cor = 0.7
T0901317 100nM vs DMSO
Cor = 0.75
log2FC in bulk RNAseq
log2FC in scRNAseq

**Supplementary Figure 17.** Correlation of gene fold changes between bulk and single cell RNA-seq for each treatment. Single cell fold changes were calculated using the mean expression across all single cells associated with a treatment compared to the mean expression across all single cells associated with DMSO. Pearson's correlations are shown.

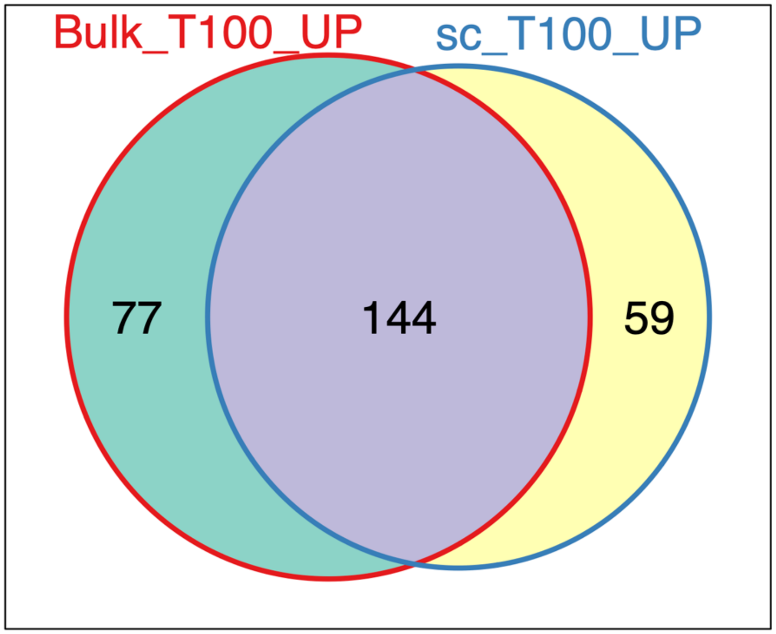

Bulk_T100_UP
sc_T100_UP
77
144
59


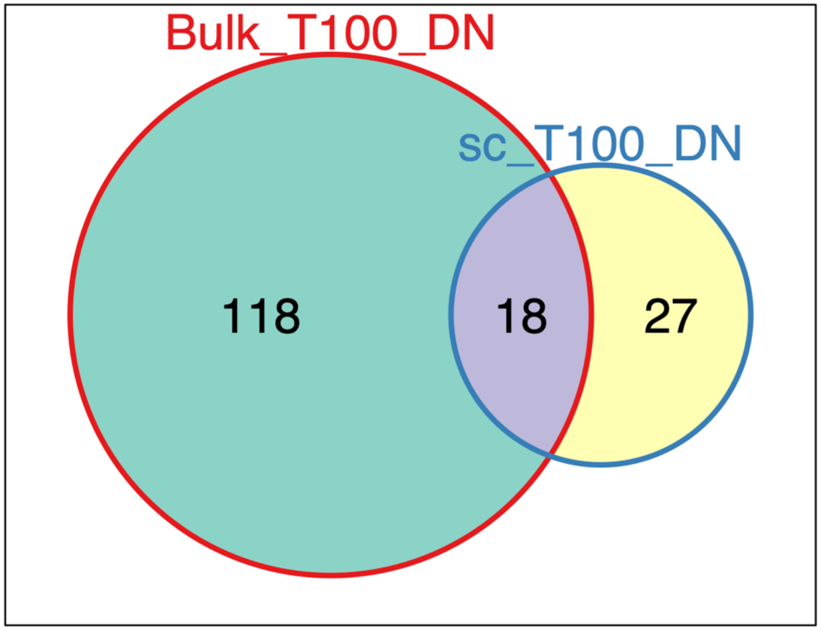

Bulk_T100_DN
sc_T100_DN
118
18
27

**Supplementary Figure 18.** Overlap in upregulated (UP) and downregulated (DN) DEGs between bulk RNA-seq and the union of single cell RNA-seq DEGs in clusters C1, C2, C3, C4, C8, and C10 with T0901317 100nM treatment. Bulk DEGs were filtered to genes tested in at least one single cell contrast.

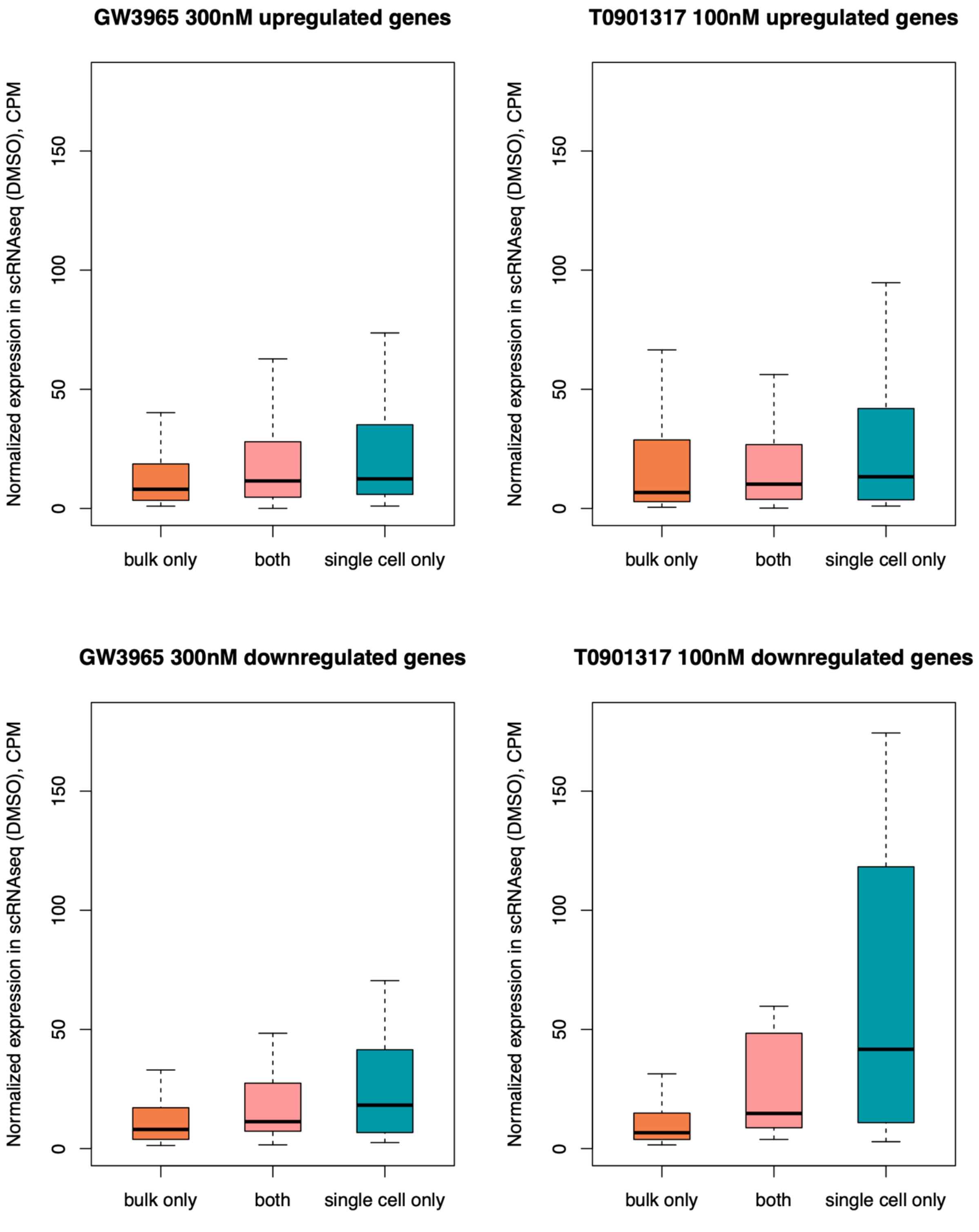
GW3965 300nM upregulated genes
T0901317 100nM upregulated genes
GW3965 300nM downregulated genes
T0901317 100nM downregulated genes
Normalized expression in scRNAseq (DMSO), CPM
0
50
100
150
bulk only
both
single cell only

**Supplementary Figure 19.** Comparison of single cell expression levels in counts per million (CPM) for DEGs identified by bulk RNA-seq only, single cell RNA-seq only, or both methods for GW3965 300nM and T0901317 100nM treatments. Bulk DEGs were filtered to genes tested in at least one single cell contrast. For each gene, its single cell expression level was calculated as the mean expression across single cells in the DMSO samples.

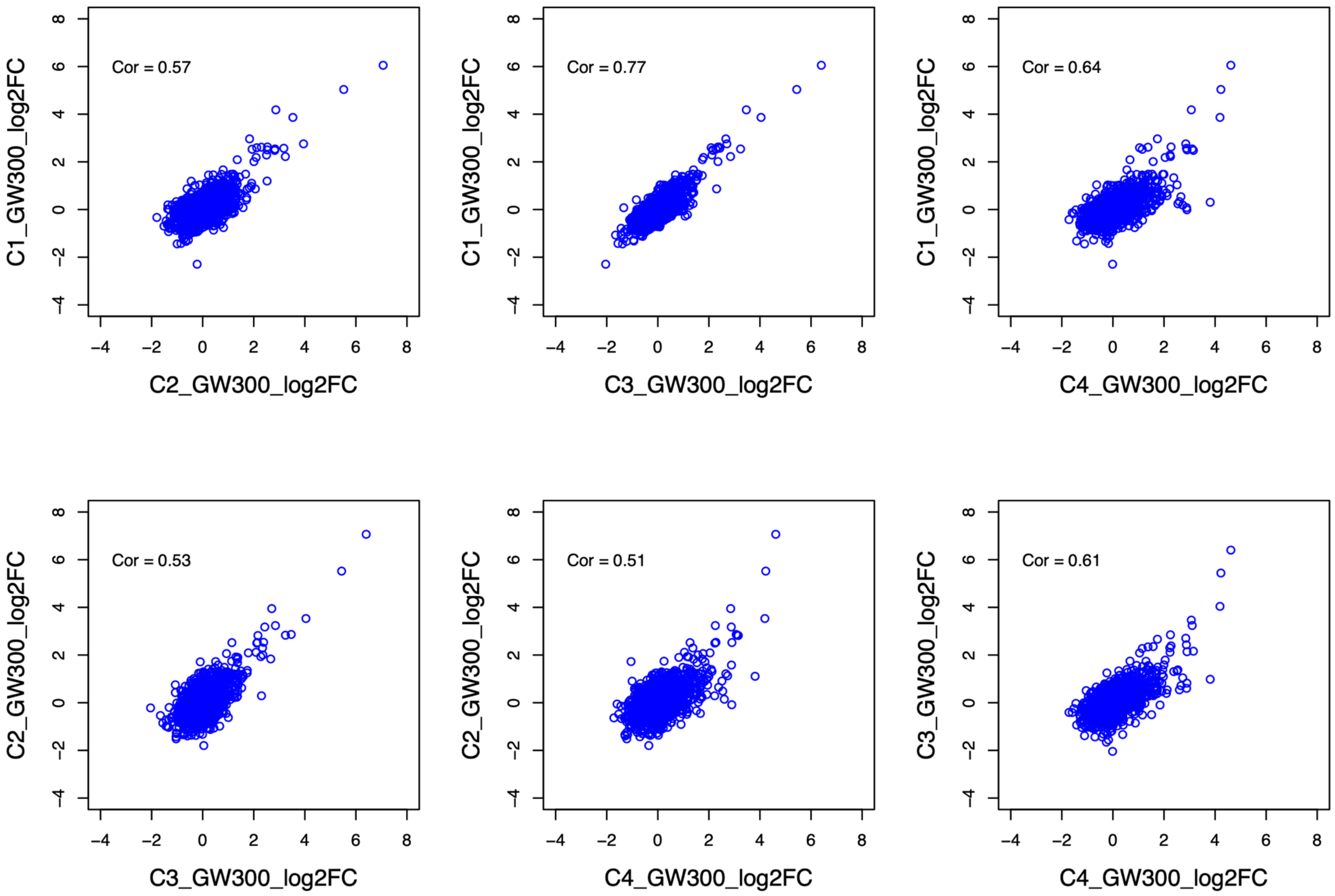

Cor = 0.57
C1_GW300_log2FC
C2_GW300_log2FC
Cor = 0.77
C1_GW300_log2FC
C3_GW300_log2FC
Cor = 0.64
C1_GW300_log2FC
C4_GW300_log2FC
Cor = 0.53
C2_GW300_log2FC
C3_GW300_log2FC
Cor = 0.51
C2_GW300_log2FC
C4_GW300_log2FC
Cor = 0.61
C3_GW300_log2FC
C4_GW300_log2FC

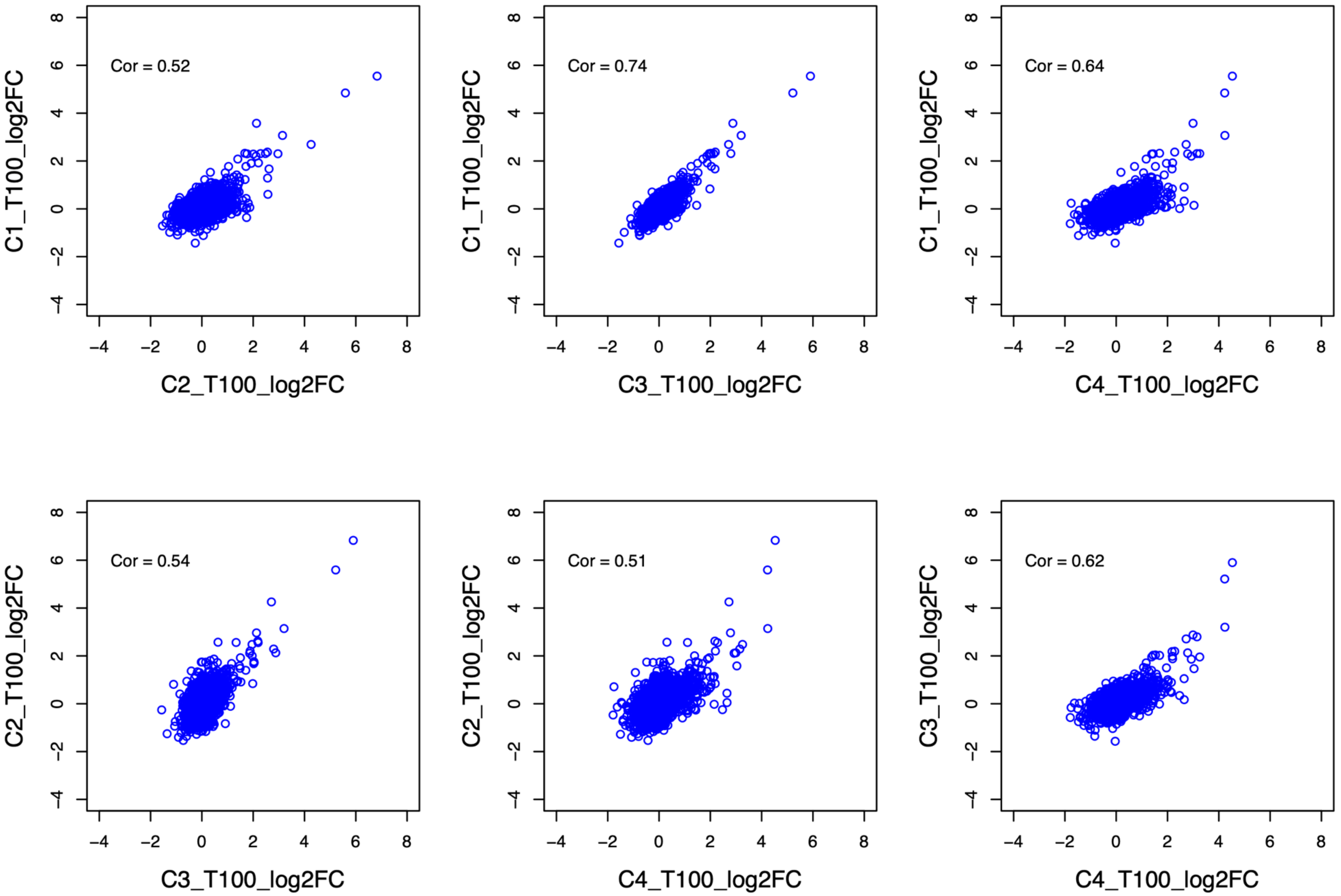

Cor = 0.52
C1_T100_log2FC
C2_T100_log2FC
Cor = 0.74
C1_T100_log2FC
C3_T100_log2FC
Cor = 0.64
C1_T100_log2FC
C4_T100_log2FC
Cor = 0.54
C2_T100_log2FC
C3_T100_log2FC
Cor = 0.51
C2_T100_log2FC
C4_T100_log2FC
Cor = 0.62
C3_T100_log2FC
C4_T100_log2FC

**Supplementary Figure 20.** Pairwise correlation of gene fold changes for the high doses of GW3965 and T0901317 across single cell clusters C1, C2, C3, and C4. Pearson's correlations are shown. GW300: GW3965 300 nM, T100: T0901317 100 nM.

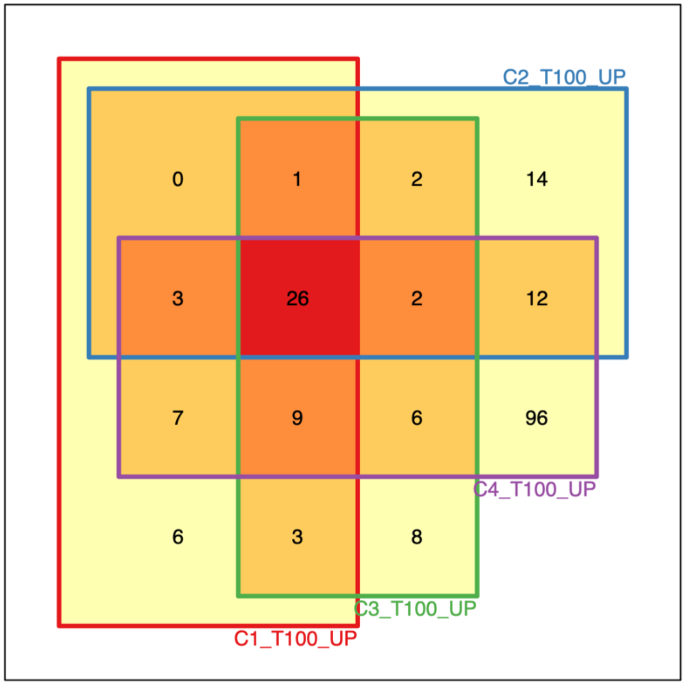
C2_T100_UP
0
1
2
14
3
26
2
12
7
9
6
96
C4_T100_UP
6
3
8
C3_T100_UP
C1_T100_UP

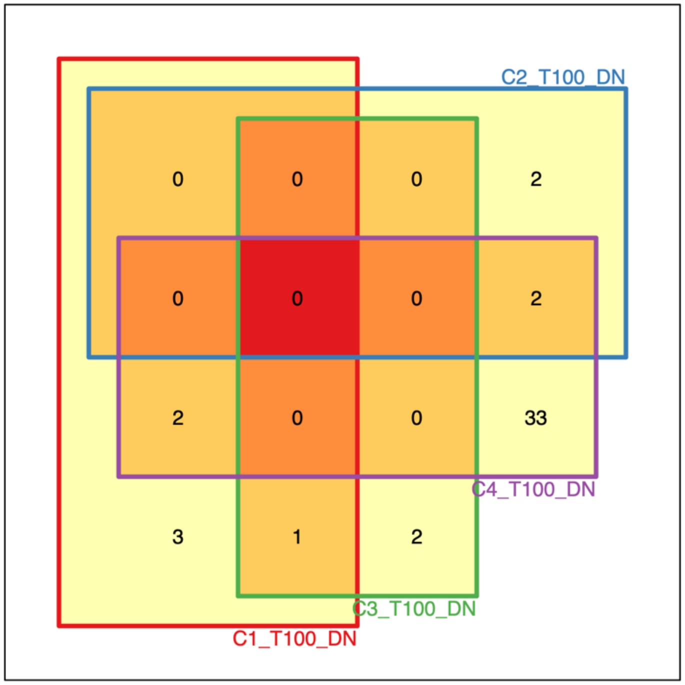
C2_T100_DN
0
0
0
2
0
0
0
2
2
0
0
33
C4_T100_DN
3
1
2
C3_T100_DN
C1_T100_DN

**Supplementary Figure 21.** Overlap in upregulated (UP) and downregulated (DN) DEGs for single cell clusters C1, C2, C3, and C4 with T0901317 100nM treatment (T100).
